# Sustaining microglial reparative function enhances stroke recovery

**DOI:** 10.1101/2024.04.10.588813

**Authors:** Jun Tsuyama, Seiichiro Sakai, Kumiko Kurabayashi, Ryuki Koyama, Yuichiro Hara, Ito Kawakami, Hideya Kawaji, Takashi Shichita

**Author notes:** Corresponding authors: Jun Tsuyama, Ph.D. and Takashi Shichita, M.D., Ph.D. 1-5-45 Yushima, Bunkyo-ku, Tokyo 113-8510, Japan.

## Abstract

Neurological symptoms after brain injury can remain as lifelong detrimental sequelae since most spontaneous brain recovery disappears within a few months after brain injury. Microglia play an essential role in recovery processes after brain injury; however, cellular and molecular mechanisms that diminish spontaneous brain functional recovery remain unknown. We discovered by cellular fate analysis that reparative myeloid cells remained in the post-stroke brain even after losing their reparative function. ZFP384 was identified as a pivotal transcriptional regulator that diminished recovery phase–associated gene expression in reparative myeloid cells, turning them into ruined cells which lost reparative functions. ZFP384 diminished the YY1-mediated chromatin interaction necessary for expressing recovery phase–associated genes. Antisense oligonucleotide against *Zfp384* sustained the broad range of neural repair effects of myeloid cells and enhanced stroke recovery, even in the chronic phase of ischemic stroke recovery. Thus, therapeutics preventing the myeloid reparative immunity from reaching a ruined state sustains brain functional recovery.

## Introduction

Brain injury, especially stroke, is a major cause of severe disability, and it shortens healthy life expectancy worldwide^1,2^. Stroke induces acute harmful inflammation and brain swelling, which worsens a patient’s prognosis; therefore, much effort is focused on acute treatments and prevention of stroke. When stroke patients overcome the acute phase, brain inflammation resolves around 1 week after stroke onset, and spontaneous brain recovery mechanisms improve the neurological deficits^3^. Rehabilitation is a common method to enhance recovery mechanisms and improve functional prognosis after brain injury; however, therapeutic drugs that enhance these recovery processes have not been developed. These spontaneous recovery processes in human patients and in stroke model animals generally disappear within a few months after brain injury^4,5^, but the cellular and molecular mechanisms that diminish the reparative functions of brain cells remain to be clarified. If brain recovery is incomplete, the remaining neurological deficits cause permanent detrimental sequelae and affect the patient for life. Thus, there is an urgent need to identify therapeutic targets that prolong the reparative function of brain cells to achieve sustainable brain recovery.

Microglia play pivotal roles in the recovery processes after brain injury. Recent studies have revealed the heterogeneity of brain myeloid cells that are essential for brain homeostasis and rapidly respond to various stresses in the brain^6-8^. Severe brain tissue injury such as stroke activates these myeloid cells to trigger inflammation and exaggerate brain damage; however, the role of these myeloid cells changes to a reparative function after inflammation resolves in the recovery phase^9,10^. Insulin-like growth factor 1 (IGF1) is a representative neurotrophic factor produced by reparative myeloid cells that promotes axonal growth and synaptogenesis^11^. The depletion of myeloid cells several days after the onset of ischemic stroke decreases IGF1 production and worsens neuronal injury^12^. IGF1-expressing microglia produce various reparative factors such as osteopontin (SPP1), neudesin neurotrophic factor (NENF), growth differentiation factors, and fibroblast growth factors, which are important for neuronal repair^13^. Thus, the roles of microglia, macrophages, and neutrophils in neuronal repair after brain injury have attracted attention as a promising therapeutic target for brain pathologies^14-16^.

To date, the cellular fate of reparative brain cells has been elusive^17^. After the reparative function of myeloid cells diminishes, they may disappear from the injured brain due to apoptotic cell death or egression. It is also possible that they remain in the injured brain as ruined or memory myeloid cells whose gene expression profiles are similar to those of homeostatic myeloid cells. Recent modalities of experimental and clinical techniques, such as comprehensive gene expression analysis and oligonucleotide therapeutics, have allowed us to investigate and manipulate the functions of microglia which mainly internalize therapeutic antisense oligonucleotide (ASO) among brain cells^18^. In this study, we discovered that the reparative myeloid cells turned into ruined cells which lost their reparative functions and remained in the post-stroke brain. Increased ZFP384 expression diminished the recovery phase-associated gene expression in reparative myeloid cells, given that myeloid-specific *Zfp384*-deficiency or intracerebroventricular administration of ASO against *Zfp384* sustained the broad range of neural repair functions of myeloid cells and enhanced stroke recovery.

## Results

### CD45^int^ CD11b^int^ cells that have lost reparative gene expression remained in the post-stroke brain

To identify the cellular fate of reparative myeloid cells, we first investigated gene expression profiles associated with the recovery phase of ischemic stroke by using *Igf1*-*Egfp* transgenic mice bearing the EGFP reporter under the promoter/enhancer region of *Igf1*, a representative marker of reparative myeloid cells ^11^. We successfully detected EGFP-expressing cells in the CD45^int^ CD11b^int^ population collected from an ischemic hemisphere but not in the contralateral normal hemisphere of post-ischemic day 6 brain (**Fig. 1a**). Most of the EGFP-expressing cells were IBA1 ^+^ in the peri-infarct region on day 6 after stroke onset (**Extended Data Fig. 1a**). The depletion of *Igf1* -expressing cells 11 days after stroke onset impaired stroke recovery (**Extended Data Fig. 1b**, **c**), demonstrating the reparative role of *Igf1*-expressing myeloid cells in the recovery phase of ischemic stroke. RNA-seq analysis of *Igf1*-expressing CD45^int^ CD11b^int^ cells in post-ischemic day 6 brain enabled us to identify 391 genes associated with the recovery phase, including *Spp1*, *Nenf*, and *Gdf15*, neurotrophic factors known to be important for neural repair (**Fig. 1b** and **Extended Data Table 1**). Gene ontologies associated with vasculature, neuron projection, and nervous system development were enriched in these 391 recovery phase–associated genes (**Fig. 1c**).

**Figure 1.**
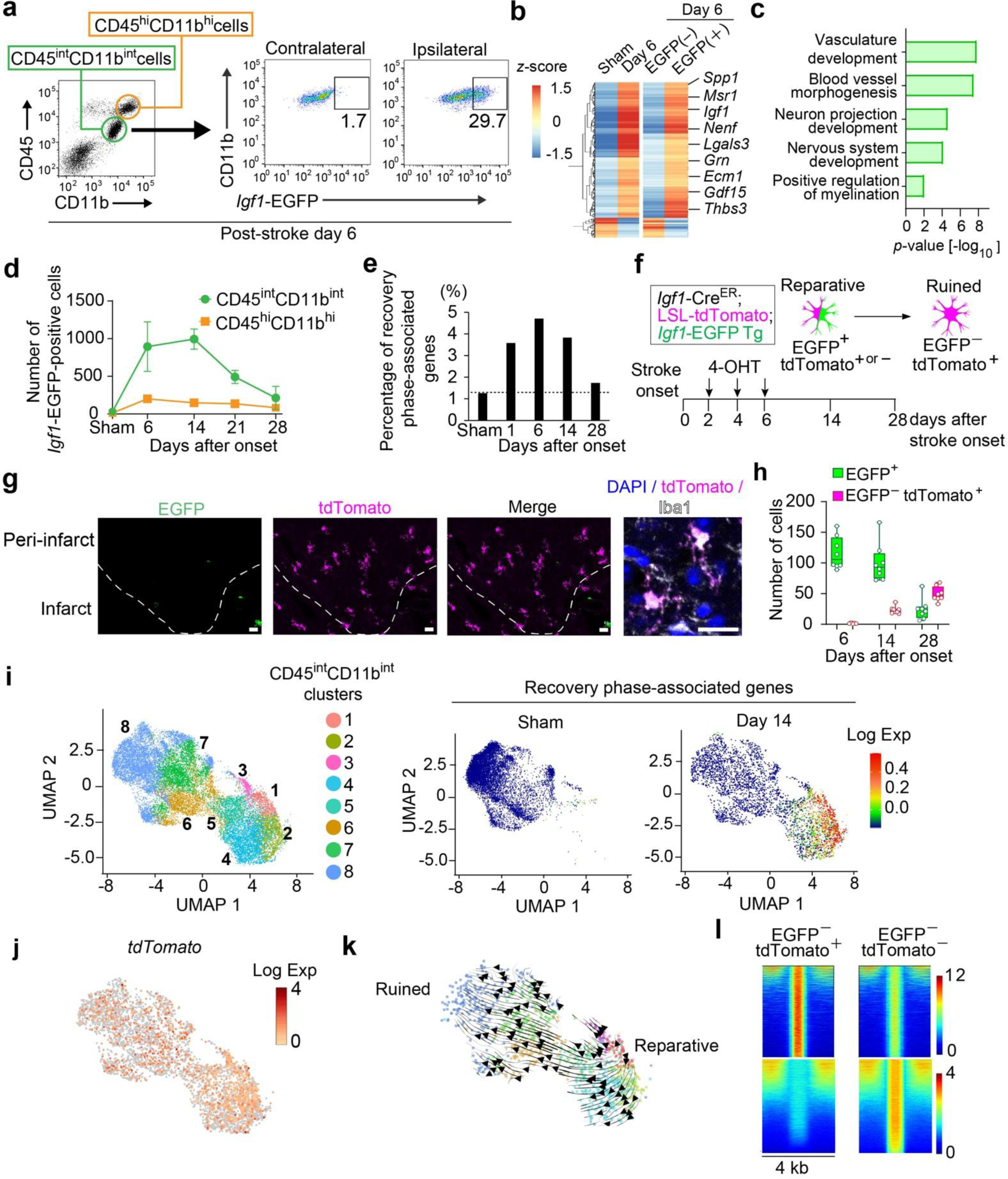
CD45^int^ CD11b^int^ cells remained in the brain even after losing recovery phase–associated gene expression. (**a**) Fluorescence-activated cell sorting (FACS) analysis of CD45^int^ CD11b^int^ cells collected from post-ischemic day 6 brain of *Igf1*-*Egfp* transgenic mice. (**b**) Heatmap of mRNA expression levels of genes showing a 2-fold increase or decrease between EGFP^–^ and EGFP^+^ CD45^int^ CD11b^int^ cells collected from sham-operated or post-ischemic day 6 brains of *Igf1*-*Egfp* transgenic mice. EGFP^+^ cells were isolated from the ipsilateral hemisphere as shown in Figure 1a. (**c**) Gene ontologies enriched in the gene list whose expression levels were increased in post-ischemic day 6 EGFP^+^ cells. (**d**) Absolute number of *Igf1*-EGFP^+^ cells in CD45^int^ CD11b^int^ or CD45^hi^CD11b^hi^ cells collected from the ipsilateral hemisphere at the indicated time point (n = 3 for each time point) after stroke onset. (**e**) Time-dependent changes in the percentage of recovery phase–associated genes in total read counts obtained from RNA-seq analysis of CD45^int^ CD11b^int^ cells collected from at least 10 mice at each time point. (**f**) The scheme explains the strategy to distinguish ruined cells (which lost recovery phase–associated gene expression) from reparative cells labeled by the 4-hydroxytamoxifen (4-OHT)-mediated *Cre* -recombination system. (**g**) Immunohistochemistry of the peri-infarct region on day 28 after stroke onset (bar: 20 μm). (**h**) Time-dependent changes in the absolute number of indicated cells (n = 8 for each) in the peri-infarct region (per 400 μm^2^). (**i**) Uniform manifold approximation and projection (UMAP) of 14,313 CD45^int^ CD11b^int^ cells (left panel) and heatmap of each sample showing recovery phase–associated gene expression level (right panel) (10,008 cells for sham; 4,305 cells for day 14). The cells were collected from sham-operated brain or the post-ischemic day 14 ipsilateral hemisphere. (**j**) Heatmap of *tdTomato* mRNA expression detected in CD45^int^ CD11b^int^ cells collected from the post-ischemic day 14 ipsilateral hemisphere. (**k**) RNA trajectory analysis of cells that expressed *tdTomato* mRNA on day 14 after ischemic stroke. Lower-right clusters are reparative cells which expressed recovery phase–associated genes as shown in Fig. 1i, and upper-left clusters are ruined cells which lost recovery phase–associated gene expression. (**l**) Difference in open-chromatin regions detected by ATAC-seq analysis between EGFP^–^tdTomato^+^ cells (ruined state) and EGFP^–^tdTomato^–^ cells (homeostatic state) collected from post-ischemic day 28 ipsilateral hemisphere.

CD45^int^ CD11b^int^ cells were the major cells expressing *Igf1*; their numbers peaked on day 14 after stroke onset but decreased thereafter (**Fig. 1d**). Coinciding with this, expression levels of the 391 recovery phase–associated genes returned to near-baseline levels on day 28 after stroke onset (**Fig. 1e**), indicating that CD45^int^ CD11b^int^ cells lost their recovery phase–associated functions around day 28 after stroke onset. To label the cells that lost their reparative roles (ruined cells), we conducted *Cre*^ER^-mediated cellular fate tracing of reparative cells that expressed *Igf1* by day 6 after stroke onset (**Fig. 1f** and **Extended Data Fig. 1d**). We discovered that cells that had lost *Igf1* expression remained in the peri-infarct region of post-ischemic day 28 brain (**Fig. 1g**). The number of these ruined cells increased from day 14 after stroke onset (**Fig. 1h**). Single-cell RNA-seq (scRNA-seq) analysis of CD45^int^ CD11b^int^ cells was used to identify the clusters (CD45^int^ CD11b^int^ clusters 1–5) bearing recovery phase–associated gene expression on day 14 after stroke onset (**Fig. 1i**). *Igf1*-*Cre^ER^*-mediated tdTomato expression was observed in all CD45^int^ CD11b^int^ clusters (**Fig. 1j**), and RNA trajectory analysis of tdTomato-positive cells demonstrated that the reparative cells bearing recovery phase–associated gene expression turned into ruined cells (**Fig. 1k**). Although the gene expression profiles of these ruined cells could not be distinguished from that of CD45^int^ CD11b^int^ cells in sham-operated mice (**Fig. 1i–k**), there were some differences between them in euchromatin regions (**Fig. 1l**). Thus, reparative CD45^int^ CD11b^int^ cells remained in the post-stroke brain even after losing recovery phase–associated gene expression, suggesting a molecular mechanism that diminished the reparative role of CD45^int^ CD11b^int^ cells.

### ZFP384 diminished recovery phase–associated gene expression in CD45^int^ CD11b^int^ cells

To identify a key transcription regulator that diminished the expression of recovery phase–associated genes, we investigated time-dependent changes in the epigenetic chromatin state in CD45^int^ CD11b^int^ cells using an assay for transposase-accessible chromatin (ATAC)-seq analysis. We found euchromatin regions specifically observed from day 14 to 28 after stroke onset (**Fig. 2a**). Transcription factor motif enrichment analysis of these euchromatin regions suggested transcription factor candidates that were functional in CD45^int^ CD11b^int^ cells from day 14 to 28 after stroke onset (**Fig. 2b**).

**Figure 2.**
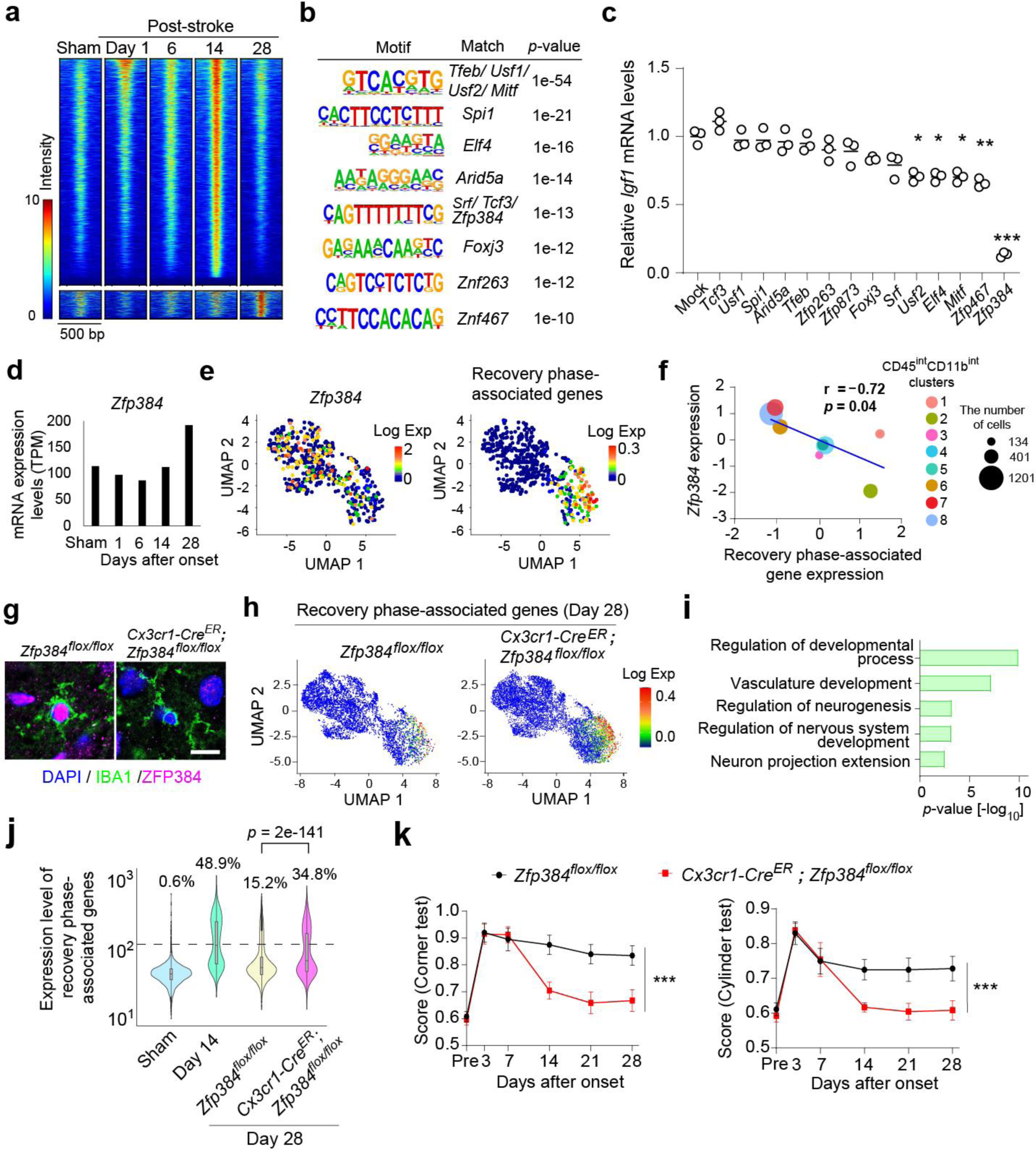
ZFP384 diminished expression of recovery phase–associated genes in CD45^int^ CD11b^int^ cells. (**a**) Open-chromatin regions detected by ATAC-seq analysis that were specifically observed in the CD45^int^ CD11b^int^ cells collected from post-ischemic day 14 or 28 ipsilateral hemisphere. (**b**) Transcription factor motif enrichment analysis of DNA sequences specifically detected by ATAC-seq in the CD45^int^ CD11b^int^ cells on day 14 or 28 after stroke onset. (**c**) Screening results of transcription factors obtained by motif enrichment analysis that diminished *Igf1* mRNA expression in the microglial cell line (n = 3 per sample). (**d**) Time-dependent change of *Zfp384* mRNA expression detected by RNA-seq analysis in the CD45^int^ CD11b^int^ cells collected from at least 10 mice after stroke onset. (**e**) Gene expression heatmaps using the UMAP of *tdTomato* -positive CD45^int^ CD11b^int^ cells (1277 cells) on day 14 after ischemic stroke. (**f**) Correlation analysis between *Zfp384* and recovery phase–associated gene mRNA expression in CD45^int^ CD11b^int^ clusters. (**g**) Immunohistochemistryof ZFP384 in peri-infarct IBA1^+^ myeloid cells on day 28 after stroke onset (bar: 10 μm). (**h**) Comparison of recovery phase–associated gene expression heatmaps of CD45^int^ CD11b^int^ cells between *Zfp384^flox/flox^* and *Cx3cr1-Cre^ER^*; *Zfp384^flox/flox^* mice on day 28 after stroke onset (5198 cells for *Zfp384^flox/flox^*; 9294 cells for *Cx3cr1-Cre^ER^*; *Zfp384^flox/flox^* mice). (**i**) Gene ontology analysis of genes whose expression levels were increased in CD45^int^ CD11b^int^ cells of *Cx3cr1-Cre^ER^*; *Zfp384^flox/flox^* mice. (**j**) Comparison of total standardized read counts of recovery phase–associated genes in all CD45^int^ CD11b^int^ cells at each time point. Percentages indicate the ratio of cells showing recovery phase–associated gene expression (>200 read counts; dashed line). (**k**) Neurological deficits after stroke onset in *Zfp384^flox/flox^* and *Cx3cr1-Cre^ER^*; *Zfp384^flox/flox^* mice (n = 10 for *Zfp384^flox/flox^*, n = 12 for *Cx3cr1-Cre^ER^*; *Zfp384^flox/flox^* mice). * *p* < 0.05, ** *p* < 0.01, *** *p* < 0.001 vs. mock (**c**) or *Zfp384^flox/flox^* mice (**j**,**k**) (one-way ANOVA with Dunnett’s test [**c**], Pearson correlation analysis [**f**], Kruskal-Wallis test with Dunn’s test [**j**], two-way ANOVA with Tukey’s test [**k**]). Error bars represent the mean ± standard error of the mean (SEM).

We found that open-chromatin regions around the *Igf1* gene locus were similar between post-stroke day 6 CD45^int^ CD11b^int^ cells and a microglial cell line that constantly expressed *Igf1* (**Extended Data Fig. 2a**). We then examined the lentiviral overexpression of the transcription factor candidates in the microglial cell line and identified *Zfp384* as a repair-terminating factor that significantly decreased *Igf1* expression (**Fig. 2c**). *Zfp384* expression levels were inversely correlated with *Igf1* expression levels (**Extended Data Fig. 2b**), and the DNA-binding ability of ZFP384 was necessary for suppression of *Igf1* expression (**Extended Data Fig. 2c**).

The expression level of *Zfp384* in CD45^int^ CD11b^int^ cells slightly decreased on day 6 but increased thereafter until day 28 after stroke onset (**Fig. 2d**), which was inversely correlated with time-dependent changes of recovery phase–associated gene expression (**Fig. 1e**). Indeed, scRNA-seq analysis revealed that *Zfp384* expression levels were higher in clusters that did not express recovery phase–associated genes (**Fig. 2e**) and inversely correlated with recovery phase–associated gene expression levels in CD45^int^ CD11b^int^ clusters on day 14 after stroke onset (**Fig. 2f**).

To clarify the role of ZFP384 in microglia and macrophages, we generated *Cx3cr1-Cre^ER^*-mediated conditional *Zfp384* -deficient mice because CD45^int^ CD11b^int^ myeloid cells included microglia and infiltrating hematopoietic macrophages after ischemic stroke that were *Cx3cr1* -positive (**Extended Data Fig. 3a**, **b**,**c**), and the expression of microglia-specific markers *Hexb* and *Sall1* was drastically decreased in CD45^int^ CD11b^int^ cells after ischemic stroke (**Extended Data Fig. 3d**). When 4-hydroxytamoxifen (4-OHT) was administered on day 7 after stroke onset, we could not detect ZFP384 expression in IBA1^+^ cells on day 28 after stroke onset (**Fig. 2g**). scRNA-seq analysis of CD45^int^ CD11b^int^ cells showed an apparent increase in expression of recovery phase–associated genes in *Cx3cr1-Cre^ER^*; *Zfp384^flox/flox^* (*Zfp384* conditional knockout [cKO]) mice compared with *Zfp384^flox/flox^* mice (**Fig. 2h** and **Extended Data Fig. 4**). Gene ontologies associated with neuron projection or developmental process of vasculature and nervous system were enriched in the genes whose expression was significantly increased in *Zfp384* cKO mice (**Fig. 2i**). The expression of recovery phase–associated genes was remarkable in CD45^int^ CD11b^int^ cells on day 14 after stroke onset and was sustained in *Zfp384* cKO mice but not in *Zfp384^flox/flox^* mice (**Fig. 2j**). Consistently, long-term neurological deficits were significantly improved in *Zfp384* cKO mice compared with *Zfp384^flox/flox^* mice (**Fig. 2k**). Infarct volume, physiological data, changes in cerebral blood flow, and survival rate were not significantly different between the two groups (**Extended Data Table 2**). These results suggest the potential of therapeutics against ZFP384 to sustain stroke recovery.

### ASO against *Zfp384* sustained stroke recovery

We next generated locked nucleic acid (LNA)-flanked ASO with various sequences against *Zfp384* mRNA (**Extended Data Fig. 5a**, **b**) and found that ASO-2 (hereafter, ASO-*Zfp384*) remarkably decreased *Zfp384* expression (**Extended Data Fig. 5c**, **d**). When ASO was conjugated with Cy3 fluorescence dye and administered into the cerebroventricle after ischemic brain injury, it was substantially internalized by CD45^int^ CD11b^int^ cells (**Fig. 3a** and **Extended Data Fig. 5e**). Compared with control ASO, ASO-*Zfp384* administration significantly decreased *Zfp384* mRNA expression in CD45^int^ CD11b^int^ cells after ischemic stroke (**Fig. 3b**).

**Figure 3.**
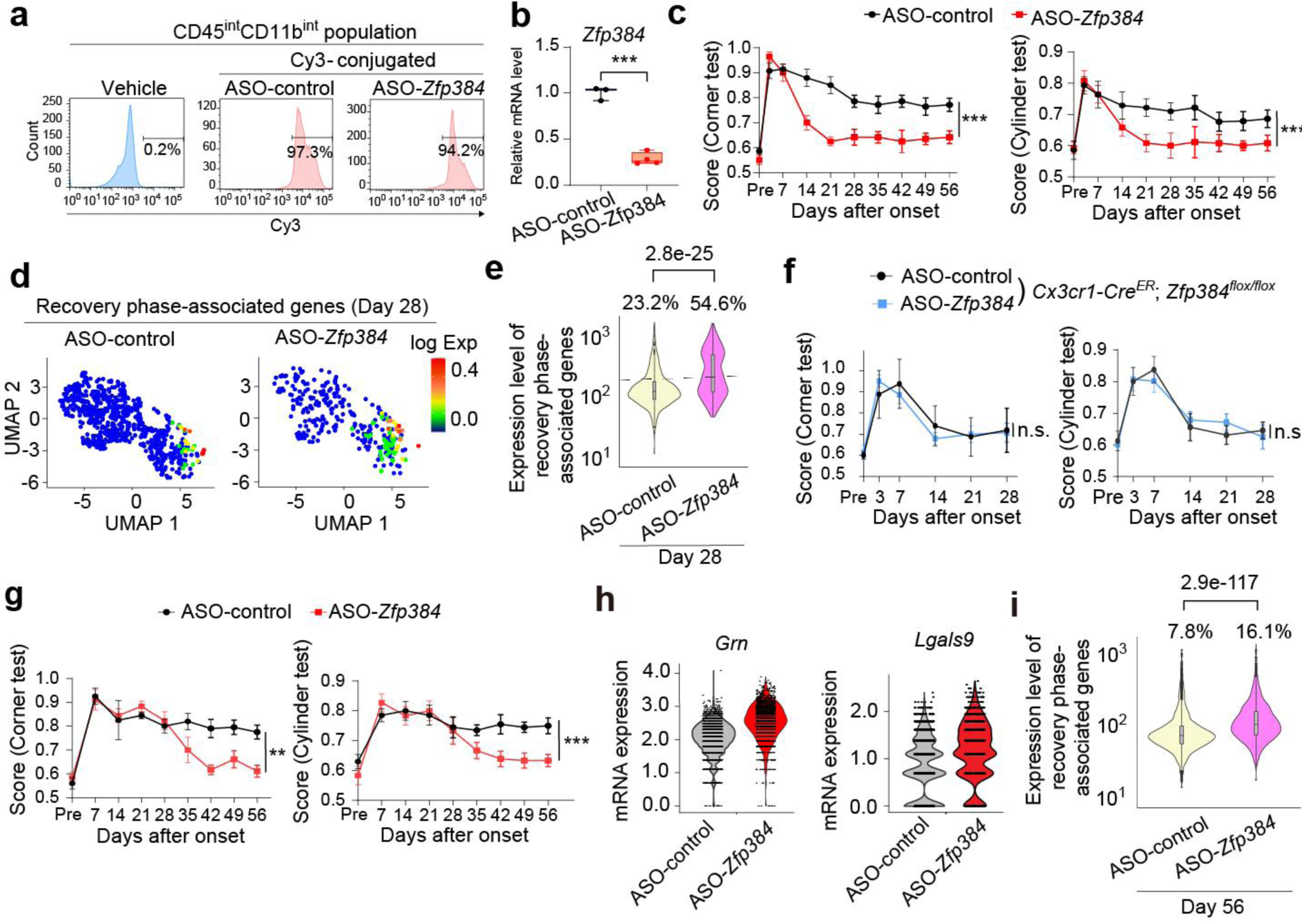
ASO against *Zfp384* prolonged the reparative function of CD45^int^ CD11b^int^ cells for sustained stroke recovery. (**a**) Internalization of fluorescent dye–conjugated ASO in CD45^int^ CD11b^int^ cells 3 days after intracerebroventricular administration. (**b**) Relative mRNA expression levels of *Zfp384* in Cy3-conjugated-ASO-internalized CD45^int^ CD11b^int^ cells collected from mice administered ASO-control or ASO-*Zfp384*. (**c**) Neurological deficits after stroke onset in mice administered ASO-control or ASO-*Zfp384* (n = 7 for ASO-control, 7 for ASO-*Zfp384*). ASOs were administered 8 and 22 days after stroke onset. (**d**) Comparison of recovery phase–associated gene expression heatmaps of CD45^int^ CD11b^int^ cells on day 28 after stroke onset between mice administered ASO-control or ASO-*Zfp384* (760 cells for ASO-control; 340 cells for ASO-*Zfp384*). (**e**) Comparison of total standardized read counts of recovery phase–associated genes in all CD45^int^ CD11b^int^ cells on day 28 after stroke onset between mice administered ASO-control or ASO-*Zfp384*. Percentages indicate the ratio of cells showing recovery phase–associated gene expression (>200 read counts; dashed line). (**f**) Neurological deficits after stroke onset in *Cx3cr1-Cre^ER^*; *Zfp384^flox/flox^* mice administered ASO-control or ASO-*Zfp384* (n = 8 for ASO-control, 7 for ASO-*Zfp384*). ASOs were administered 8 and 22 days after stroke onset. (**g**) Neurological deficits after stroke onset in mice administered ASO-control or ASO-*Zfp384*(n = 10 for ASO-control, 9 for ASO-*Zfp384*). ASOs were administered 29 and 43 days after stroke onset. (**h**) Comparison of reparative gene expression in all CD45^int^ CD11b^int^ cells on day 56 after stroke onset between mice administered ASO-control or ASO-*Zfp384* (3569 cells for ASO-control; 1992 cells for ASO-*Zfp384*). (**i**) Comparison of total standardized read counts of recovery phase–associated genes in all CD45^int^ CD11b^int^ cells on day 56 after stroke onset between mice administered ASO-control or ASO-*Zfp384*. Percentages indicate the ratio of cells showing recovery phase–associated gene expression (>200 read counts; dashed line). \*\**p* < 0.01, *** *p* < 0.001 vs. ASO-control (**b**,**c**, **e**,**f**,**g**, **i**) (two-sided Student’s *t*-test [**b**], two-way ANOVA with Tukey’s test one-way ANOVA with Dunnett’s test [**c**, **f**,**g**], Mann-Whitney U test [**e**,**i**]). n.s.: not significant. Error bars represent the mean ± standard error of the mean (SEM).

The long-term neurological deficits after ischemic stroke were significantly improved when ASO-*Zfp384* was administered on day 8 after stroke onset (**Fig. 3c**). Infarct volume, changes in cerebral blood flow, and survival rate were not significantly different between ASO-control- and ASO-*Zfp384* -administered mice (**Extended Data Table 3**). A similar therapeutic effect of ASO-*Zfp384* was observed in aged mice (**Extended Data Fig. 6a**). ASO-*Zfp384* increased the expression of recovery phase–associated genes (**Fig. 3d** and **Extended Data Fig. 4**) and significantly increased the ratio of CD45^int^ CD11b^int^ populations that expressed recovery phase–associated genes on day 28 after stroke onset (**Fig. 3e**). ASO-*Zfp384* administration did not affect the CD45^hi^CD11b^hi^ populations (**Extended Data Fig. 6b**, **c**) and did not improve long-term neurological deficits in *Zfp384* cKO mice (**Fig. 3f**), indicating that ASO-*Zfp384* exerted its therapeutic effects through CD45^int^ CD11b^int^ cells.

Furthermore, even when administered on day 29 after stroke onset, ASO-*Zfp384* improved neurological deficits (**Fig. 3g**). ASO-*Zfp384* administration on day 29 after stroke onset sustained the expression of recovery phase–associated genes such as *Grn* and *Lgals9* (**Fig. 3h**) and increased the ratio of reparative CD45^int^ CD11b^int^ populations on day 56 after stroke onset (**Fig. 3i** and **Extended Data Fig. 4**). Thus, ASO against *Zfp384* sustained functional recovery, even in the chronic phase of ischemic stroke, by prolonging the reparative roles of CD45^int^ CD11b^int^ cells.

### ASO against *Zfp384* promoted neural repair after ischemic stroke

To confirm the enhanced neural repair achieved by sustaining the reparative function of CD45^int^ CD11b^int^ cells, we compared the single-cell RNA-seq results of brain cells on day 28 after stroke onset between mice administered ASO-control and ASO-*Zfp384* (**Fig. 4a**). The highest numbers of differentially expressed genes (DEGs) were detected in oligodendrocyte precursor cells (OPCs) and excitatory neurons (**Fig. 4b**). In these glial and neuronal cells, ontologies associated with a broad range of neural repair, such as neuronal differentiation, neuron projection development, and synaptic plasticity, were enriched in DEGs whose expression levels increased with ASO-*Zfp384* administration (**Fig. 4c–e**).

**Figure 4.**
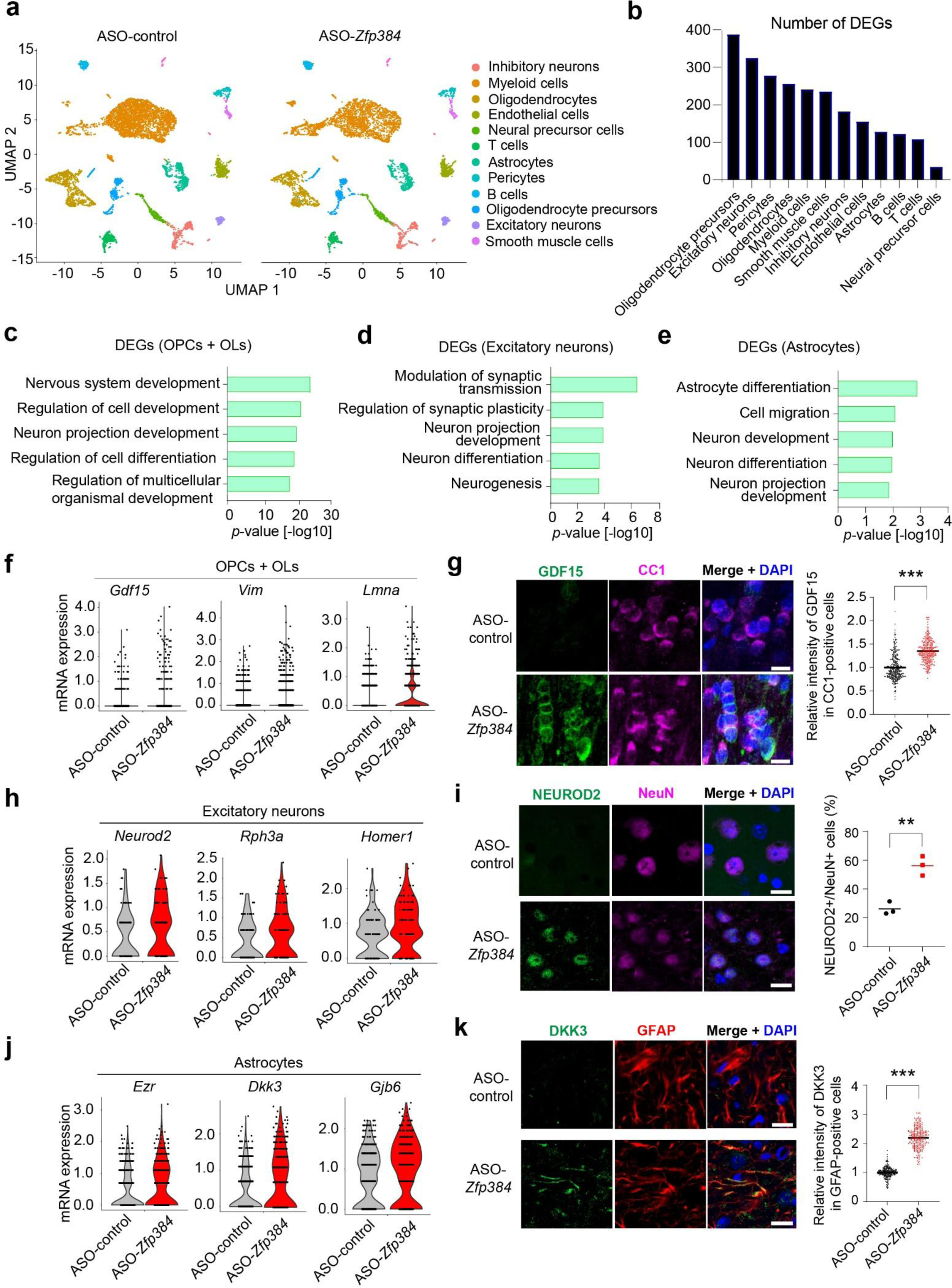
ASO against *Zfp384* had broad-range neural repair effects in the chronic phase of ischemic stroke. (**a**) UMAP of brain cells collected from post-ischemic day 28 ipsilateral brain hemisphere of mice administered ASO-control or ASO-*Zfp384*(10,008 cells for ASO-control; 4305 cells for ASO-*Zfp384*). (**b**) Number of differentially expressed genes (DEGs) in each cellular population when compared between mice administered ASO-control or ASO-*Zfp384*. (**c**, **d**,**e**) Results of gene ontology analysis of DEGs whose expression increased in mice administered ASO-*Zfp384*. OPCs: oligodendrocyte precursor cells, OLs: oligodendrocytes. (**f**,**h**,**j**) Violin plots of representative DEGs in each cell type. (**g**, **i**,**k**) Immunohistochemistry and mean fluorescence intensity of GDF15, NEUROD2, and DKK3 in each cell type. ** *p* < 0.01, *** *p* < 0.001 vs. ASO-control (**g**, **i**,**k**) (two-sided Student’s *t*-test [**g**, **i**,**k**]).

The mRNA expression levels of *Gdf15*, *Vim*, and *Lmna*, important factors for myelinating function and axonal regeneration ^19-21^, were increased in mice administered ASO-*Zfp384* (**Fig. 4f**). Increased expression of GDF15 in oligodendrocytes (OLs) was confirmed by immunohistochemistry of ischemic brain tissue 28 days after stroke onset (**Fig. 4g**). ASO-*Zfp384* administration increased mRNA expression of *Neurod2*, *Rph3a*, and *Homer1* in excitatory neurons, which promote synaptic maturation, dendritic spine growth, and functional recovery from neurological deficits ^22-24^ (**Fig. 4h**). The ratio of NEUROD2-positive neurons in the peri-infarct region was increased by ASO-*Zfp384* administration (**Fig. 4i**). Expression levels of *Ezr*, *Dkk3*, and *Gjb6* were increased in astrocytes of mice administered ASO-*Zfp384* (**Fig. 4j**); these genes encode proteins that are important for functional improvement following stroke and synaptic plasticity ^25-27^. Immunohistochemistry of ischemic brain tissues on day 28 after stroke onset revealed increased expression of DKK3 in peri-infarct astrocytes (**Fig. 4k**). Thus, ASO against *Zfp384* exerted a broad range of neural repair effects in neurons and glial cells by sustaining myeloid reparative function.

### ZFP384 diminished YY1-mediated recovery phase–associated gene expression in CD45^int^CD11b^int^ cells

We attempted to clarify the molecular mechanisms underlying the ZFP384-mediated loss of reparative function in CD45^int^ CD11b^int^ cells. HiChIP analysis, which detects chromatin loops maintaining genomic interactions among enhancer and promoter regions (E-P interactions), revealed that chromatin loops were apparently formed around the *Igf1*locus in CD45^int^ CD11b^int^ cells 6 days after stroke onset, compared to sham-operated mice (**Fig. 5a**). Indeed, the number of chromatin loops associated with E-P interactions significantly increased in the recovery phase–associated genes 6 days after stroke onset (**Fig. 5b**), implying the involvement of a chromatin looping factor in recovery phase–associated gene expression after ischemic stroke.

**Figure 5.**
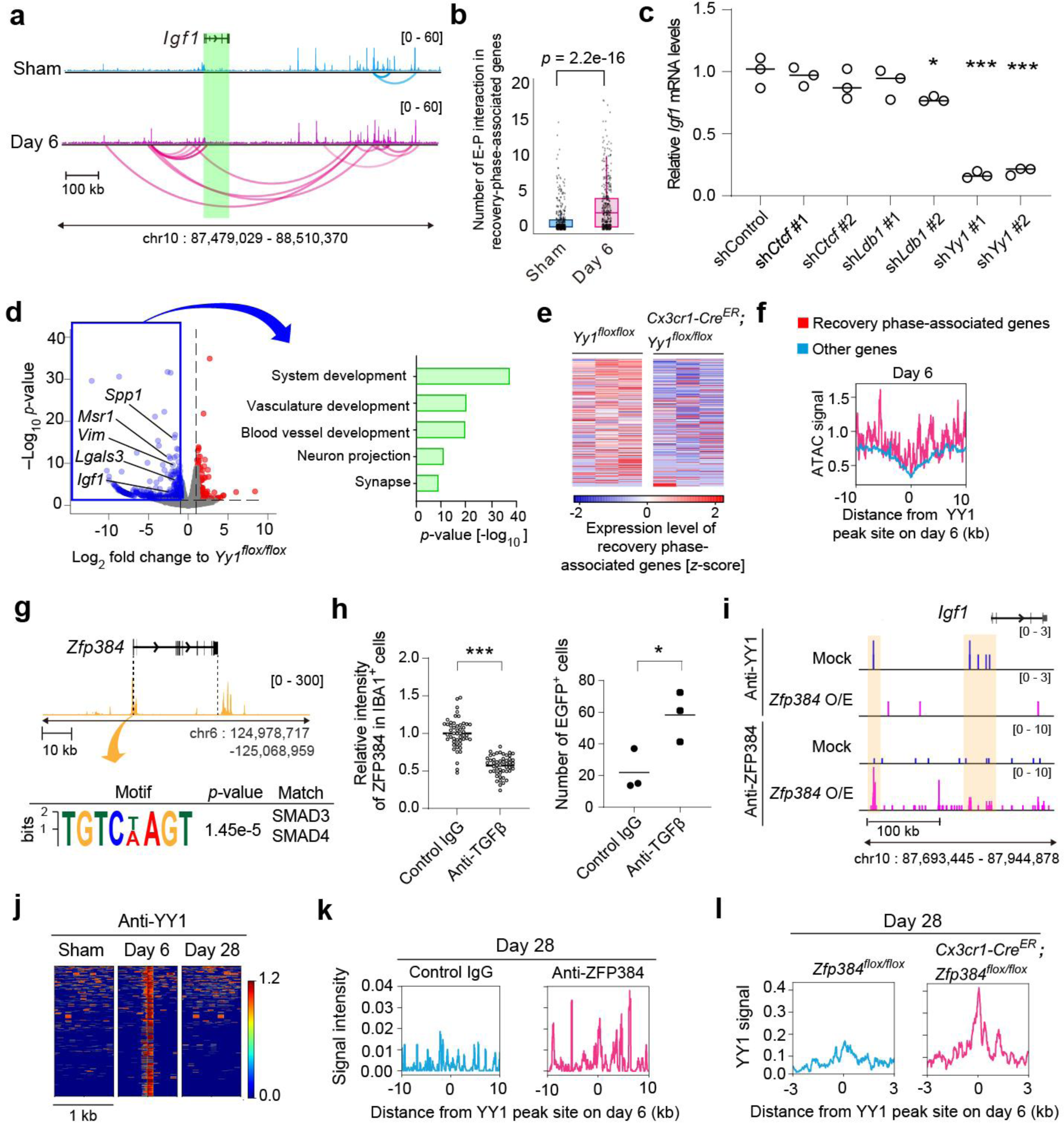
ZFP384 disrupted YY1-mediated recovery phase–associated gene expression in CD45^int^CD11b^int^ cells. (**a**) Results of chromatin loop detection around the *Igf1* locus by HiChIP in CD45^int^ CD11b^int^ cells before and 6 days after ischemic stroke onset. The peaks indicate the intensities of ATAC-seq signals. (**b**) Comparison of enhancer-promoter (E-P) interactions around recovery phase–associated genes before and 6 days after stroke onset. (**c**) Relative *Igf1* mRNA expression levels in each short-hairpin-RNA-introduced (shRNA-introduced) microglial cell line. #1 and #2 indicate different shRNA sequences against *Ctcf*, *Ldb1*, or *Yy1*. (**d**) RNA-seq results of post-ischemic day 6 CD45^int^ CD11b^int^ cells in *Cx3cr1-Cre^ER^*; *Yy1^flox/flox^* and *Yy1^flox/flox^* mice. Right panel shows the gene ontology analysis of the genes whose expression levels significantly decreased in *Cx3cr1-Cre^ER^*; *Yy1^flox/flox^* mice compared to *Yy1^flox/flox^* mice. (**e**) Heatmaps show the comparison of recovery phase–associated gene expression levels between *Yy1^flox/flox^* and *Cx3cr1-Cre^ER^*; *Yy1^flox/flox^* mice. (**f**) Comparison of ATAC-seq signal intensities around the YY1-bound region between recovery phase–associated gene loci and other gene loci in post-ischemic day 6 CD45^int^ CD11b^int^ cells. (**g**) Upper panel shows the ATAC-seq signals detected around *Zfp384* gene in post-ischemic day 28 CD45^int^ CD11b^int^ cells; lower panel shows the result of transcription factor motif enrichment analysis of DNA sequences detected by ATAC-seq analysis in the promoter region of *Zfp384* gene. (**h**) Relative intensity of ZFP384 in IBA1^+^ cells or the number of EGFP^+^IBA1^+^ cells in the peri-infarct region analyzed by immunohistochemistry on day 28 after stroke onset. Control IgG or anti-TGF β antibody was administered on day 14 and day 21 after stroke onset. (**i**) Results of cleavage under targets and tagmentation (CUT&Tag) analysis using anti-YY1 or anti-ZFP384 antibody around *Igf1* gene in a mock-transduced or *Zfp384* -overexpressing (O/E) microglial cell line. (**j**) YY1-bound regions observed in post-ischemic day 6 CD45^int^ CD11b^int^ cells were compared to sham-operated or post-ischemic day 28 CD45^int^ CD11b^int^ cells. (**k**) ZFP384-binding signal intensities in post-ischemic day 28 CD45^int^ CD11b^int^ cells around the YY1-bound regions detected in post-ischemic day 6 CD45^int^ CD11b^int^ cells. (**l**) Comparison of YY1-binding signal intensities in post-ischemic day 28 CD45^int^ CD11b^int^ cells between *Zfp384^flox/flox^* and *Cx3cr1-Cre^ER^*; *Zfp384^flox/flox^* mice around the YY1-bound regions detected in post-ischemic day 6 CD45^int^ CD11b^int^ cells. * *p* < 0.05, \*\*\**p* < 0.001 vs. sham (**b**), shControl (**c**), or control IgG (**h**) (Mann-Whitney U test [**b**], one-way ANOVA with Dunnett’s test [**c**], two-sided Student’s *t*-test [**h**]). Error bars represent the mean ± standard error of the mean (SEM).

We then examined knockdown of chromatin looping factors, such as CTCF, LDB1, and YY1, in the microglial cell line and found that YY1 was important for *Igf1* expression (**Fig. 5c**). Microglia/macrophage-specific deficiency of *Yy1* significantly decreased gene expression associated with vasculature development and neuron projection in CD45^int^ CD11b^int^ cells 6 days after stroke onset (**Fig. 5d**). *Yy1* deficiency impaired the recovery phase–associated gene expression in CD45^int^ CD11b^int^ cells (**Fig. 5e**). Consistent with this, cleavage under targets and tagmentation (CUT&Tag) analysis demonstrated a euchromatin state around YY1-bound regions at recovery phase–associated gene loci compared with other gene loci (**Fig. 5f**).

ATAC-seq analysis of CD45^int^ CD11b^int^ cells on day 28 after stroke onset revealed that the binding motif of SMAD3 or SMAD4, transcription components of TGF β signaling, was enriched in the promoter region of *Zfp384* gene (**Fig. 5g**). TGF β appeared to increase ZFP384 expression and impair the reparative function of IBA1^+^ cells, given that administration of anti-TGF β antibody decreased ZFP384 levels but increased the number of EGFP^+^ cells in *Igf1*-*Egfp* transgenic mice (**Fig. 5h**).

We discovered that *Zfp384* overexpression in the microglial cell line removed YY1 from the promoter and enhancer regions of *Igf1* gene; instead, overexpressed ZFP384 bound to these regions (**Fig. 5i**). In the murine model of ischemic stroke, YY1-bound regions seen in CD45^int^ CD11b^int^ cells on day 6 were rarely detected on day 28 after stroke onset (**Fig. 5j**). Around these YY1-bound regions that were observed on day 6, binding of ZFP384 was detected on day 28 (**Fig. 5k**). Finally, we examined *Cx3cr1-Cre^ER^*; *Zfp384^flox/flox^* mice and found that binding of YY1 was maintained until day 28 in the YY1-bound regions observed in the CD45^int^ CD11b^int^ cells on day 6 compared with that in *Zfp384^flox/flox^* mice (**Fig. 5l**). These results indicated that increased ZFP384 removed YY1 from the transcriptional complex necessary for expression of recovery phase–associated genes.

### ZFP384 expression was inversely correlated with IGF1 expression in human post-stroke brain

We examined human brain tissue to investigate the correlation between ZFP384 (ZNF384 in human) and IGF1 after ischemic stroke. In the peri-infarct region on day 6 after stroke onset, we found that IGF1^+^ IBA1^+^ myeloid cells were ZNF384^+^ (**Fig. 6a**). Although expression of ZNF384 was not detected in IBA1^+^ cells of normal brain obtained from control patients, IGF1^–^ ZNF384^+^ cells were observed on post-stroke day 49 in the peri-infarct region (**Fig. 6a**). IGF1^+^ IBA1^+^ cells were observed in the peri-infarct region until 3 weeks after stroke onset but rarely observed thereafter (**Fig. 6b**). In contrast, the number of ZNF384^+^ IBA1^+^ cells increased from 2 weeks after stroke onset and these cells were present in the peri-infarct regions of post-stroke brain from week 5 to 7 (**Fig. 6c**). The number of ZNF384^+^ cells was inversely correlated with the number of IGF1^+^ cells in a time-dependent manner after ischemic stroke onset (**Fig. 6d**), indicating the clinical relevance of therapeutic strategies inhibiting ZFP384 for sustaining brain functional recovery.

**Figure 6.**
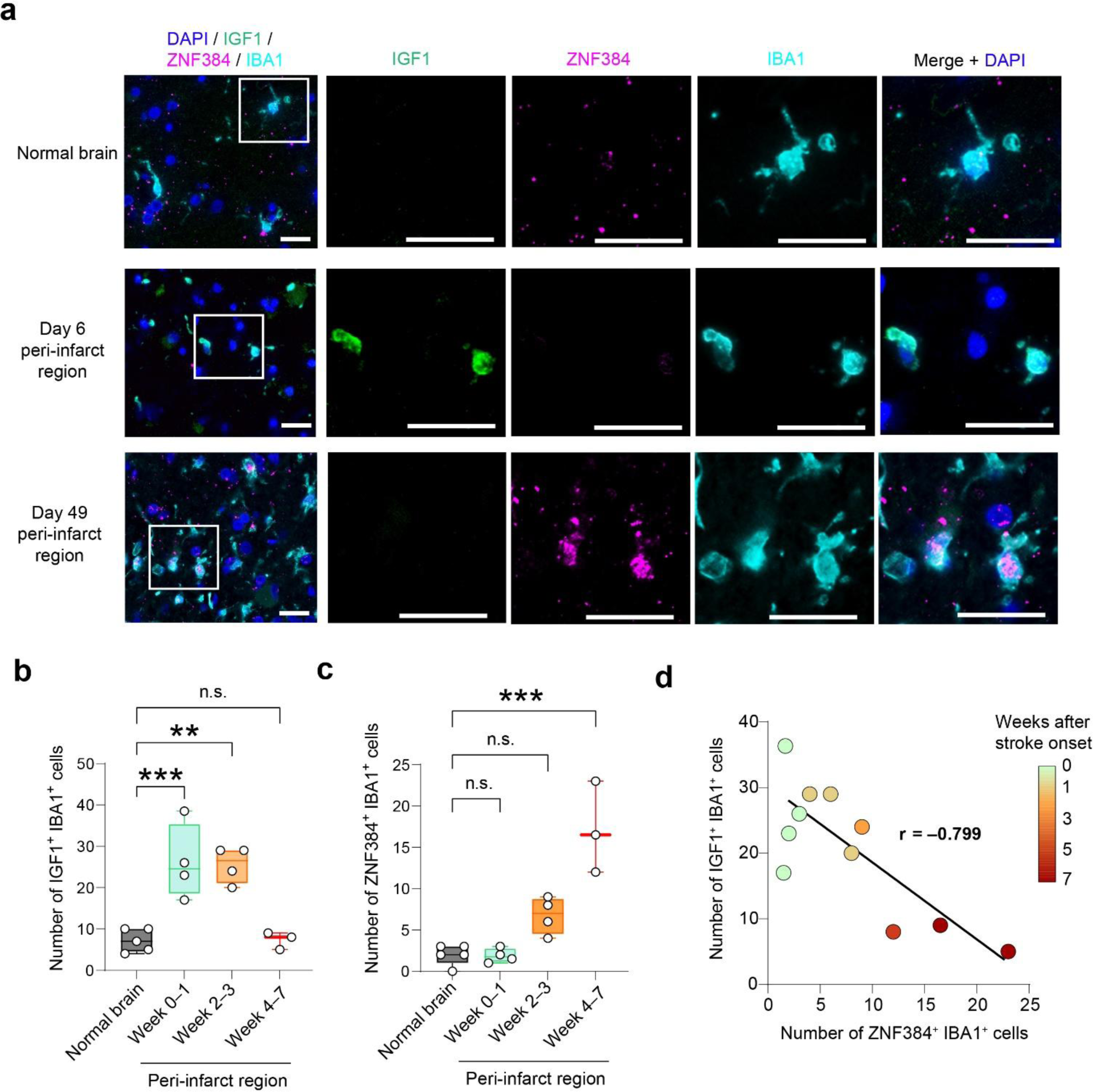
ZFP384 expression is inversely correlated with IGF1 expression in the peri-infarct region of human post-stroke brain. (**a**) Immunohistochemistry of ZFP384, IGF1, and IBA1 in normal brain of control patients without ischemic stroke (5 cases) and the peri-infarct region of ischemic stroke patients (11 cases). Scale bar: 10 μm. (**b**,**c**) Time-dependent changes of IGF1 expression (**b**) and ZNF384 expression (**c**) in the peri-infarct IBA1^+^ cells of ischemic stroke patients. (**d**) Correlation analysis between the number of ZNF384^+^IBA1^+^ and IGF1^+^IBA1^+^ cells in the peri-infarct region of ischemic stroke patients. Heatmap shows the time elapsed since stroke onset. ** *p* < 0.01, \*\*\**p* < 0.001 vs. normal brain (**b**,**c**) (one-way ANOVA with Dunnett’s test [**b**,**c**], Pearson’s correlation analysis [**d**]). n.s.: not significant.

## Discussion

Myeloid cells have important roles in tissue repair after the inflammatory phase of tissue injury. Microglial cells (i.e., CD45^int^ CD11b^int^ cells), which partly include blood-derived myeloid cells in the case of brain injuries ^28^, play important roles in brain functional recovery. In our study, reparative microglial cells, consisting of several clusters, showed a broad range of reparative effects on neurons and glial cells. This explains the remarkable functional recovery associated with ASO-*Zfp384* administration on day 8 after stroke onset, since infarct volume does not change much after the first week following stroke onset. Chronic exposure to TGF β, which is enriched in brain tissue, increased ZFP384 expression and impaired microglial reparative immunity. Although microglial ZFP384 expression was inversely correlated with recovery phase–associated gene expression in human ischemic stroke, it is possible that similar molecular mechanisms underlying ruined reparative immunity also work in other brain pathologies or organ injuries.

ZFP384 and YY1 are nuclear matrix proteins (NMPs) that comprise dynamic scaffold for intranuclear reactions and intrachromosomal interactions ^29^, although the importance of interaction among NMPs has not been clarified (summarized figure: **Extended Data Fig. 7**). Because YY1 (NMP1) is a multifunctional transcriptional regulator that consists of chromatin loops, it was considered a key functional component of the transcriptional complex driving microglial recovery phase–associated gene expression after ischemic stroke. On the other hand, the DNA-binding activity of ZFP384 (NMP4) is necessary for suppressing *Igf1* mRNA expression (**Extended Data Fig. 2c**). In the terminal phase of stroke recovery, increased ZFP384 bound to recovery phase–associated gene loci to remove YY1 from the transcriptional complexes, diminishing expression of recovery phase–associated genes. CUT&Tag analysis detected the other chromatin regions to which YY1 and ZFP384 were adjacently bound, suggesting that the interactions between YY1 and ZFP384 may be different in various transcriptional regulator complexes.

ASO-mediated inhibition of *Zfp384* sustained the expression of recovery phase–associated genes in microglial cells without increasing expression of inflammatory mediators such as cytokines or chemokines. The ASO dose used in the current study did not result in any neurological symptoms in mice, and we did not observe any harmful effects on brain cells by sustaining microglial reparative function in single-cell transcriptomic studies of post-stroke brain cells. ASO can be administered intrathecally at the lumbar level to achieve delivery to the cerebroventricle in humans. Because ASO was detectable in microglial cells for at least 2 weeks after administration, ASO against *Zfp384* could enhance long-term rehabilitation effects in stroke patients with minimal invasiveness. Our results illustrate a mechanism to allow the development of therapeutics to aid in long-term functional recovery of the brain following stroke.

## Acknowledgments

We thank Mrs. Yoshiko Yogiashi for their experimental assistance throughout the study and Dr. Yuichi Hiraoka for advice to generate *Igf1*-*Cre^ER^* mice and *Zfp384^flox/flox^* mice by gene editing. We appreciate Dr. Hiroshi Shitara and Dr. Rie Ishii for supporting animal breeding, Mrs. Mariko Yoshimura for assistance with writing this manuscript, and Dr. Erika Seki for the kind cooperation in preparing human brain sections. We also appreciate kind support from Prof. Tomoyuki Furuyashiki and Prof. Shuh Narumiya for supplying the *Cx3cr1-Cre^ER^* mice. This work was supported by CREST from AMED under grant number JP23gm1210010 (T.S.); a Grant-in-Aid for Scientific Research on Innovative Areas (Dynamic Regulation of Brain Function by the Glia decoding [23H04181]) from the Ministry of Education, Culture, Sports, Science and Technology of Japan (MEXT) (T.S.); JSPS KAKENHI Grants-in-Aid for Scientific Research (B) (21H02820) (T.S.); Scientific Research (C) (21K06386) (S.S.) and (23K05626) (J.T.); a Toray Science and Technology Grant (T.S.); the Takeda Science Foundation (T.S., S.S., J.T.); the Uehara Memorial Foundation (S.S.); and grants from the MSD Life Science Foundation (T.S.), Senri Life Science Foundation (T.S.), Ono Pharmaceutical Foundation for Oncology, Immunology, and Neurology (T.S.); Charitable Trust Mihara Cerebrovascular Disorder Research Promotion Fund (T.S.); Medical Research Center Initiative for High Depth Omics, TMDU, Nanken-Kyoten, TMDU, Multilayered Stress Diseases (JPMXP1323015483), TMDU.

## Author contributions

J.T. designed and performed the experiments, analyzed the data, and wrote the manuscript; S.S., K.K., and R.K. performed the experiments and provided technical advice; Y.H. and H.K. analyzed the RNA-seq data and participated in discussion, I.K. provided the samples and advice in the analysis of human stroke patients; T.S. initiated and directed the entire study, designed the experiments, and wrote the manuscript.

## Competing financial interests

The authors declare no competing financial interests.

## Materials and Methods

### Mice

All mice were maintained in a conventional facility at the Tokyo Medical and Dental University or Tokyo Metropolitan Institute of Medical Science in Tokyo, Japan. All experiments were approved by the Institutional Animal Research Committee of the Tokyo Medical and Dental University (approval number: A2023-172C2) and Tokyo Metropolitan Institute of Medical Science (approval number: 21-004). C57BL/6J mice were used for WT mice. *Igf1*-*Egfp* mice (MMRRC 000261-UNC) were crossed with C57BL/6J mice several times before being used in experiments. Rosa26-CAG-LSL-tdTomato (JAX Stock No: 007908) or ROSA26-iDTR mice (JAX Stock No: 007900) were bred with *Igf1*-IRES-*Cre ^ER^* mice (C57BL/6J background) several times and used for experiments. *Cx3cr1* -*Cre* ^ERT2^ mice (*Cx3cr1-Cre ^ER^*: JAX Stock No: 021160) were bred with *Zfp384 ^flox/flox^* mice (C57BL/6J background) several times and used for experiments. Littermates were used for the experiment using *Yy1 ^flox/flox^* mice (JAX Stock No: 014649) mice.

### Generation of gene-edited mice

*Igf1*-IRES-*Cre ^ER^* mice and *Zfp384 ^flox/flox^* mice were generated by microinjection of crRNA (Integrated DNA Technologies), tracrRNA (Integrated DNA Technologies), donor plasmid DNA, and Cas9 protein into zygotes. To generate *Igf1-Cre ^ER^* mice, a gRNA sequence targeting the 3 ′ UTR of the *Igf1* gene locus was used (gRNA sequence: 5 ′-CAAAGGAUCCUGCGGUGAUG-3 ′). The donor plasmid was constructed with the IRES2 sequence and ERT2-*Cre* -ERT2 fusion protein inserted between homology arms of 1.5–2 kb each. To generate *Zfp384 ^flox/flox^* mice, gRNA sequences targeting intron 8 of *Zfp384* (NM_001372424) were used (gRNA sequence: 5′-CAUUUAUAACUGGUGCCCUG-3 ′). A loxP-flanked genomic sequence of approximately 6.7 kb, containing exons 7 to 9 and centered on the gRNA target region, was cloned into a donor plasmid, and the protospacer adjacent motif (PAM) sequence was removed from the donor plasmid to prevent cleavage by Cas9.

### Murine model of ischemic stroke

Experiments were conducted on male mice between 8 and 24 weeks of age, with a weight range of 20 to 30 g. Male and female mice between 19 and 24 months were used for the aged mice experiment. Random selection was used for all experiments. A total of 8 to 15 mice was deemed necessary to achieve adequate statistical power for comparing neurological deficits. Transient brain ischemia, induced by middle cerebral artery occlusion (MCAO), was generated in the mice. Anesthesia was induced by 3% isoflurane with a mixture of 70% nitrous oxide and 30% oxygen, and maintained by 1% isoflurane during the operation. Silicone-coated monofilaments were then inserted through the common carotid artery to occlude the origin of the middle cerebral artery (MCA). Reperfusion was facilitated by withdrawing the inserted monofilaments 60 minutes after MCAO. Throughout the MCAO procedure, the head temperature was maintained at 36°C. Inclusion criteria for the study required confirmation of a >60% reduction in cerebral blood flow using laser Doppler flowmetry. Supplementary tables present information on the changes in cerebral blood flow during ischemia, the number of excluded mice, physiological data, and the survival rate. To evaluate the neurological deficits, we conducted the corner and cylinder tests (details of the experimental methods have been provided elsewhere; ^3^). For the cylinder test, the neurological score was calculated as follows: (number of non-impaired forelimb contacts – number of impaired forelimb contacts)/(number of non-impaired forelimb contacts + number of impaired forelimb contacts). For the corner test, the neurological score was calculated as follows: (number of ipsilateral turns)/(number of all turns).

### Depletion of Igf1-expressing brain cells

*Igf1*-IRES-*Cre*; *Igf1*-*Egfp*; ROSA26-iDTR or *Igf1*-*Egfp*; ROSA26-iDTR mice were intraperitoneally injected with 4-hydroxytamoxifen (4-OHT; H6278, Sigma) at 10 mg kg ^−1^ daily from day 4 to day 14 after stroke onset. Diphtheria toxin (BioAcademia Inc.) was injected into the cerebroventricle at 500 ng kg ^−1^ on days 11 and 14 after stroke onset. The optimal dose of diphtheria toxin was determined by assessing the depletion efficiency of *Igf1*-expressing brain cells by flow cytometry and immunostaining.

### Isolation of the immune cell-enriched population

Sham-operated mice or mice after MCAO were transcardially perfused with saline, and then the forebrain was removed. After homogenization with RPMI-1640, 1 mg ml ^–1^ collagenase (Sigma-Aldrich) and 50 μg ml ^–1^ DNase I (Sigma-Aldrich) were added to the homogenates. Brain homogenates were digested for 25 min and subjected to Percoll (GE Healthcare) gradient centrifugation. From the interlayer between 37% and 70% Percoll, cells were isolated and stained with anti-CD45 FITC (30-F11, eBioscience), anti-CD11b PerCP-Cy5.5 (M1/70, eBioscience), and anti-CD31 PE/Cyanine7 (390, eBioscience). For *Igf1* -EGFP mice, anti-CD45 APC (30-F11, eBioscience) was used. The cells were washed with PBS and analyzed using a FACSAria III cell sorter (BD/Thermo Fisher Scientific).

To examine mRNA expression levels, total RNA was purified using a ReliaPrep RNA Cell Miniprep System (Promega) and applied to quantitative PCR analysis. First-strand cDNA was synthesized using a PrimeScript RT Master Mix (RR036, Takara). Polymerase chain reactions were carried out using a SsoFast EvaGreen Supermixes (Bio-Rad) on the CFX Connect (Bio-Rad).

### RNA-seq

Total RNA was purified using a RNeasy micro kit (Qiagen) and used for the preparation of RNA-seq libraries by an Ovation SoLo RNA-Seq Library Preparation Kit (Tecan), according to the manufacturer’s procedure. Following the assessment of quantity and fragment size distribution of cDNA libraries by Agilent TapeStation 4150, the libraries were sequenced by Illumina HiSeq. Reads were aligned to the mm10 annotation file using STAR, and the read counts for each gene were determined through featureCounts. Volcano plots of gene expression levels were depicted using the replicated samples from three independent experiments after normalization, and DEGs were identified by edgeR. Genes exhibiting significant increases or decreases in expression between the two groups were subjected to gene ontology analysis using DAVID (https://david.ncifcrf.gov/).

### ATAC-seq

An ATAC-seq library was prepared according to the protocol described elsewhere ^3^. For the fragmentation and tagmentation of genomic DNA, 50,000 nuclei isolated from CD45 ^int^CD11b ^int^ cells were treated with Tn5 transposase (Illumina, Nextera DNA Library Preparation Kit). The DNA fragments were amplified using KAPA HiFi HS ReadyMix (Kapa Biosystems) and purified using SPRI select beads (Beckman). Upon assessing the quantity and fragment size distribution of the ATAC-seq libraries with Agilent TapeStation 4150, libraries were sequenced using the Illumina HiSeq sequencer, and reads were mapped to mm10 (genome file) by Bowtie2. ATAC-seq signal peaks were detected by MACS2 default parameters.

### Transcription factor motif enrichment analysis

De novo motif analysis was performed on accessible chromatin peaks obtained by ATAC-seq analysis. Motif enrichment was performed using the HOMER function findMotifsGenome. The output report links de novo motifs to known motifs and gives a motif matching score; known motifs that matched de novo motifs with a score >0.7 were linked to transcription factors based on the transcription factor catalog. MEME (https://meme-suite.org/meme/) was used to identify transcription factors that bind to the chromatin accessible regions within 1 kb of the *Zfp384* promoter region in CD45 ^int^CD11b ^int^ cells.

### Lentiviral transduction

BV2, a microglial cell line, was supplied by Prof. Kazuhide Inoue at Kyushu University, Japan. cDNA was cloned into the lentiviral vector CSII-EF-MCS-P2A-mCherry modified from CSII-EF-MCS-IRES2-Venus (RIKEN). cDNA sequences were obtained from the CCDS database (NCBI). To generate recombinant lentivirus, cDNA expression vectors with VSV-G expression vector (pCMV-VSV-G-RSV-Rev, RIKEN) and packaging vector (pMDLg/p-RRE) were transfected using polyethylenimine (PEI) into HEK293 cells supplied by the RIKEN Bioresource Center (RCB2202). Forty-eight hours after the vector-containing culture medium was replaced with fresh medium, lentivirus-containing culture medium was concentrated by centrifugation and resuspended in RPMI-1640.

### Construction and administration of ASOs

ASOs were designed to induce endogenous RNaseH-mediated degradation by complementarily binding to *Zfp384* mRNA. ASO-control which were previously used elsewhere ^30^ had no targeted genes and its sequence was GGCcaatacgccgTCA (lower case: phosphorothioate-modified DNA, upper case: phosphorothioate-modified LNA). In the ASOs against *Zfp384*, all internucleotide linkages were phosphorothioates, three bases at both ends were LNA modified, and the all the cytosine residue in the LNA was 5-methylcytosine. These ASOs were synthesized by Ajinomoto BioPharma, Inc. To examine the effects of ASOs, BV2 cells overexpressing *Zfp384* -P2A-mCherry via lentivirus were treated with ASO-control or ASOs against *Zfp384* at a concentration of 2 μM. Seventy-two hours after ASO treatment, the fluorescence intensity of mCherry in BV2 cells was analyzed by FACS to evaluate the suppression efficiency of ASOs.

ASOs (5 nmol) dissolved in PBS were administered intracerebroventricularly on days 8 and 22 after stroke onset or on days 29 and 43 after stroke onset. To investigate the uptake efficiency of ASO in brain cells, 1 nmol of Cy3-conjugated ASOs was administered intracerebroventricularly on day 1 after stroke onset. Forty-eight hours after administration, fluorescence intensity in the cells isolated from the ischemic brain was analyzed by FACS.

### scRNA-seq analysis

Brain cells including neurons were isolated from adult ischemic brain according to the protocol established in the Allen Institute ^31^; this detailed experimental method was described previously ^32^. Single CD45 ^int^CD11b ^int^ cells or brain cells were loaded onto chromium chips with a capture target of 2,000 to 10,000 cells per sample. Libraries of these samples were prepared using Chromium Next GEM Single Cell 5 ′ Reagent Kits v2 (10x Genomics) for CD45 ^int^CD11b ^int^ cells or Chromium Next GEM Single-cell 3 ′ Reagent Kits v3.1 (10x Genomics) for brain cells, according to the manufacturer’s protocol. On a DNBSEQ-G400 sequencer (MGI), obtained libraries were sequenced with a targeted sequencing depth of more than 50,000 read pairs per cell. Read mapping and counting were processed with the 10x Genomics Cell Ranger 6.0.1 with the mm10 reference genome assembly with the addition of dTomato RNA sequences and gene annotation provided by 10x Genomics ^33^. The SC Transformation method incorporated in Seurat 4.3.0 was used for the normalization of spliced-read counts for the individual genes ^34,35^. The single-cell gene expression profiles of the samples were integrated using the anchor-based integration method in Seurat ^36^. Ten groups with k-means were obtained by this integrated count matrix, which clustered the cells by the shared nearest neighbor approach implemented in Seurat. The UMAP plots were generated with the Seurat Dimplot module by using the normalized spliced read counts.

The expression of 391 recovery phase–associated genes was integrated with AddModuleScore and visualized with FeaturePlot. The read counts of recovery phase–associated genes were shown in Violinplot after normalization by the SC Transformation method. The total read counts of recovery phase–associated genes were averaged in each cell cluster and presented in bubble plots of *z*-scores for comparison among myeloid clusters. The velocyto 0.17.17 package from the BAM files generated by Cell Ranger was used for the computation of spliced and unspliced read counts, and the ratio of these read abundances for the individual cells was computed by total read numbers of these two conditions in a cell.

### Immunohistochemistry

Mice were euthanized with deep sedation and then transcardially perfused with PBS followed by 4% paraformaldehyde. Forebrains were removed and fixed with 4% paraformaldehyde overnight at 4°C, cryoprotected in sucrose, and embedded in O.C.T. compound (Sakura Finetechnical Co. Ltd.). Sections (14-to 30-μm thick) were made by cryostat and processed for immunohistochemistry. The samples were rinsed with PBS and pretreated with PBS containing 0.1% Triton-X100 for 15 min at room temperature. The sections were boiled in citrate buffer using a microwave oven. After blocking with Blocking One Histo (Nacalai Tesque), the sections were incubated with anti-IBA1 (Fuji Film, 013-27691), anti-ZFP384 (Sigma, HPA004051), anti-GFAP (Invitrogen, 13-0300), anti-GFP (Aves, GFP-1010), anti-NEUROD2 (abcam, ab104430), anti-APC (also known as CC1; Millipore, OP80), anti-DKK3 (Proteintech, 10365-1-AP), anti-NeuN (Millipore, MAB377), or anti-GDF15 (Proteintech, 27455-1-AP) overnight at 4°C. Secondary antibodies, anti-rabbit IgG antibody conjugated with Alexa Fluor 488 (Invitrogen, A11070) or Alexa Fluor 546 (Invitrogen, A11071), anti-mouse IgG antibody conjugated with Alexa Fluor 488 (Invitrogen, A11017) or Alexa Fluor 546 (Invitrogen, A11018), and anti-chicken IgY antibody conjugated with Alexa Fluor 488 (Invitrogen, A-11039) were used for detection of the primary antibody. Images of the sections were captured using a fluorescence microscope (BZ-X710, Keyence) or a confocal laser microscope (LSM980, Carl Zeiss).

For immunohistochemistry of human brain tissue obtained from ischemic stroke patients and control brain samples from individuals without ischemic stroke, the sections were deparaffinized and boiled in citrate buffer using a microwave oven. Sections were pretreated with PBS containing 0.1% Triton-X100 for 15 min and incubated with TrueBlack (Biotium) to suppress autofluorescence. After blocking with Blocking One Histo (Nacalai Tesque), the sections were incubated with anti-IGF1 (Origene, TA805748), anti-ZNF384 (Sigma, HPA004051), and anti-IBA1 (Abcam, ab40657) antibody overnight at 4°C. Anti-rabbit IgG antibody conjugated with Alexa Fluor 488 (Invitrogen, A11070) or Alexa Fluor 546 (Invitrogen, A11071), anti-mouse IgG antibody conjugated with Alexa Fluor 488 (Invitrogen, A11017) or Alexa Fluor 546 (Invitrogen, A11018), and anti-rat IgG antibody conjugated with Alexa Fluor 647 (Invitrogen, A78947) were used as secondary antibodies to detect the primary antibody. After the sections were treated with TrueVIEW Autofluorescence Quenching Kit (Vector Laboratories), images of the sections were captured using a fluorescence microscope (BZ-X710, Keyence). The absolute numbers of ZNF384- or IGF1-positive IBA1 ^+^ cells within a 0.4-mm ^2^ area in normal brain region or peri-infarct region were averaged to evaluate each section.

### CUT&Tag analysis

BV2 cells or CD45 ^int^CD11b ^int^ cells (100,000–200,000 cells from the sham-operated or ischemic hemisphere) were pelleted for 3 min at 600× *g* at room temperature and used to prepare the CUT&Tag library using a bench top CUT& Tag V.2 protocol with slight modifications. The detailed experimental procedure was described previously ^3^. Anti-YY1 (Abcam, ab109237) and anti-ZNF384 (Sigma, HPA004051) antibodies were used for primary antibodies. Sequencing was performed using the PE150 strategy (HiSeq 2500, Illumina). Reads were mapped by the same process as used for ATAC-seq. Duplicate reads were removed with picard-tools-1.119 and peak called with SEACR. Peaks with only one unique read count were then removed and plotted by deepTools.

### HiChIP analysis

CD45 ^int^CD11b ^int^ cells (200,000–300,000 cells from the sham-operated or ischemic hemisphere) were analyzed by HiChIP according to the protocol described elsewhere. After cells were sonicated for 2 min using a Covaris S220 ultrasonicator (Covaris), 2 μg of antibody against H3K27ac (Abcam, ab4729) and 20 μl of protein A beads (Thermo Fisher) were added to capture the chromatin-antibody complex. Captured chromatin was removed from protein A beads and purified by DNA Clean & Concentrator (Zymo, D4013) to be quantified by Qubit 3.0 (Thermo Fisher). After treatment with Tn5 transposase (Illumina, Nextera DNA Library Preparation Kit) for fragmentation and tagging, the library was prepared by amplification PCR (14 cycles) followed by the purification using SPRI select.

The peak file was prepared by expanding ATAC-seq signals to 1 kb that were obtained and merged from ATAC-seq analysis of CD45 ^int^CD11b ^int^ cells collected from sham-operated and post-ischemic day 6 mice. HiChIP paired-end reads were aligned to the mm10 genome using the HiC-Pro 3.1.0 pipeline of default settings ^37^. To analyze the loops connecting chromatin accessibility regions (enhancer-promoter [E-P] interactions), significant loops were called from the peak file and the validPairs file filtered by HiC-Pro using FitHiCHIP(S) (bin size: 10,000, low distance threshold: 10,000, high distance threshold: 1,000,000) ^38^. To visualize the HiCHIP contacts, we used WashU Epigenome Browser.

### Statistical analysis

Data are displayed as minimum to maximum box-and-whisker plots or means ± the standard error of the mean (SEM) in each figure. The differences between two groups were analyzed by unpaired Student’s t-test or Mann–Whitney U test. To analyze the differences among three or more groups, statistical significance was determined by analysis of variance (ANOVA) followed by post hoc multiple comparison tests (Dunnett’s test, Tukey’s test, or Sidak test) or Kruskal-Wallis test followed by post hoc multiple comparison tests (Dunn’s test). p < 0.05 was considered to represent a statistically significant difference.

## Notes

### Competing Interest Statement

The authors have declared no competing interest.

