## Extended Data Figure 1-7, Extended Data Table 1-3 for "Sustaining microglial reparative function enhances stroke recovery"

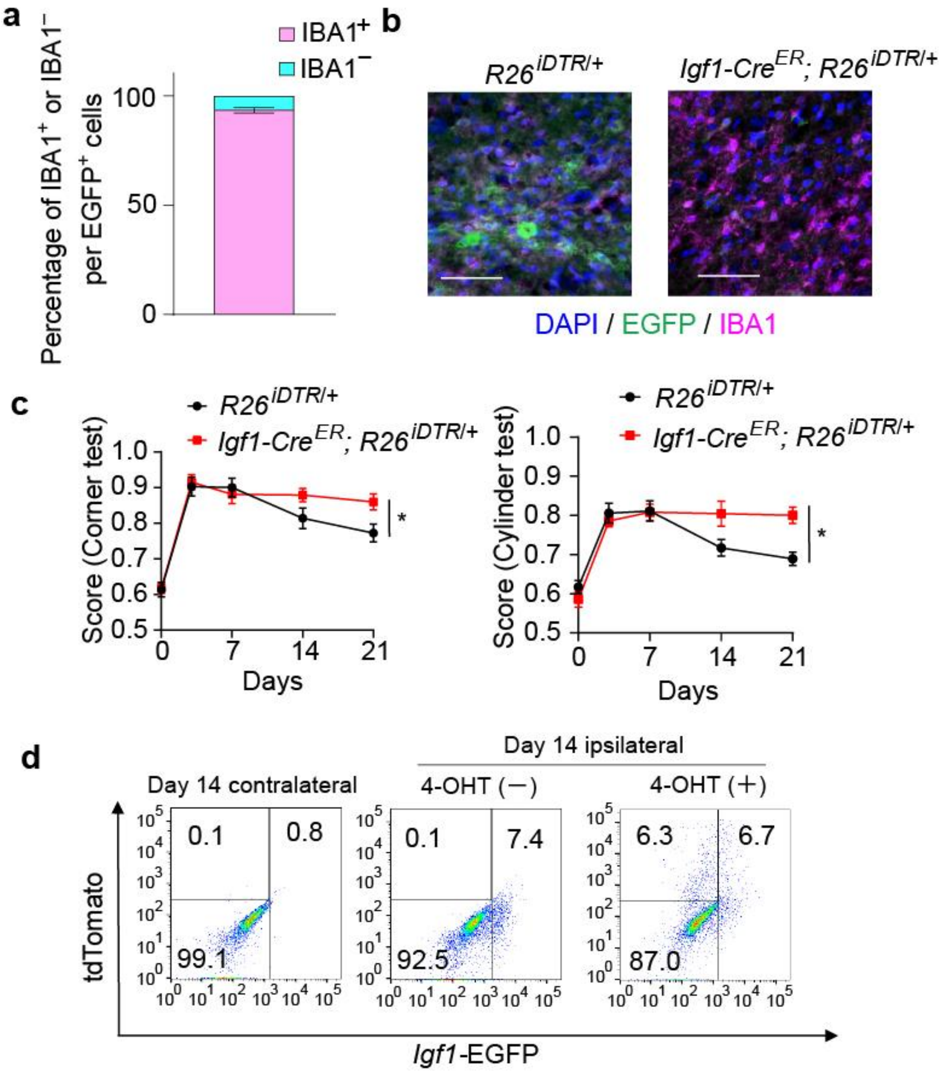


**Extended Data Figure 1. Analysis of IGF1-positive cells in the post-ischemic brain.** (**a**) Percentage of EGFP$^{+}$IBA1$^{+}$ cells and EGFP$^{+}$IBA1$^{-}$ cells in the peri-infarct region of *Igf1-Egfp* transgenic mice on day 6 after stroke onset. (**b**) Confirmation of EGFP$^{+}$ cell deletion by intracerebroventricular diphtheria toxin (DT) administration (on days 11 and 14) in *Igf1-Cre*$^{ER}$; Rosa26-loxP-stop-loxP-DTR; *Igf1-Egfp* Tg mice on day 21 after stroke onset. (**c**) Neurological deficits after stroke onset in DT-administered R26$^{iDTR}$ or *Igf1-Cre*$^{ER}$; R26$^{iDTR}$ mice (n = 10 for R26$^{iDTR}$, 12 for *Igf1-Cre*$^{ER}$; R26$^{iDTR}$ mice). (**d**) Fluorescence-activated cell sorting (FACS) analysis of post-ischemic day 14 CD45$^{int}$CD11b$^{int}$ cells collected from *Igf1-Cre*$^{ER}$; Rosa26-loxP-stop-loxP-*tdTomato*; *Igf1-Egfp* Tg mice. 4-hydroxytamoxifen (4-OHT) was administered on days 2, 4, and 6 after stroke onset. *$p$ < 0.05 vs. R26$^{iDTR}$ mice (**c**) (two-way ANOVA with Tukey's test [**c**]). Error bars represent the mean ± standard error of the mean (SEM).

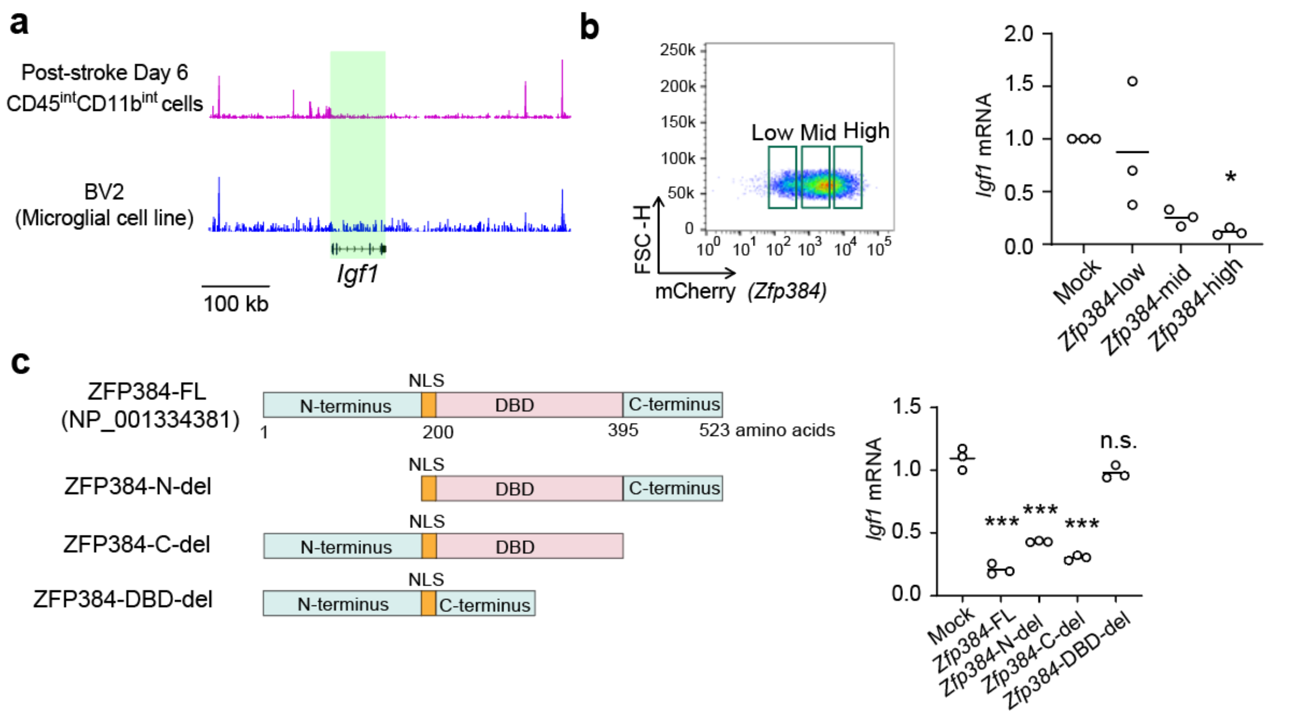


**Extended Data Figure 2. Analysis of *Zfp384* function in *Igf1* expression in microglial cells.** (**a**) Results of ATAC-seq analysis of *Igf1* gene locus in microglial cell line and post-ischemic day 6 CD45$^{int}$CD11b$^{int}$ cells. (**b**) Relative *Igf1* mRNA expression levels in mock-transduced or *Zfp384*-overexpressing microglial cell line. *Zfp384*-overexpressing cells were divided into 3 groups by *Zfp384*-P2A-*mCherry* expression levels. (**c**) Left panel shows the construct of mutant ZFP384 proteins. Numerals under the bar indicate amino acid residue numbers of ZFP384 protein. DBD: DNA-binding domain, NLS: nuclear localization signal, FL: full-length. Right panel shows the relative *Igf1* mRNA expression levels in a mock-transduced microglial cell line or cells that overexpressed the full-length or mutant *Zfp384*. *$p < 0.05$, ***$p < 0.001$ vs. mock (**b**,**c**) (one-way ANOVA with Dunnett's test [**b**,**c**]). Error bars represent the mean ± standard error of the mean (SEM). n.s.: not significant.

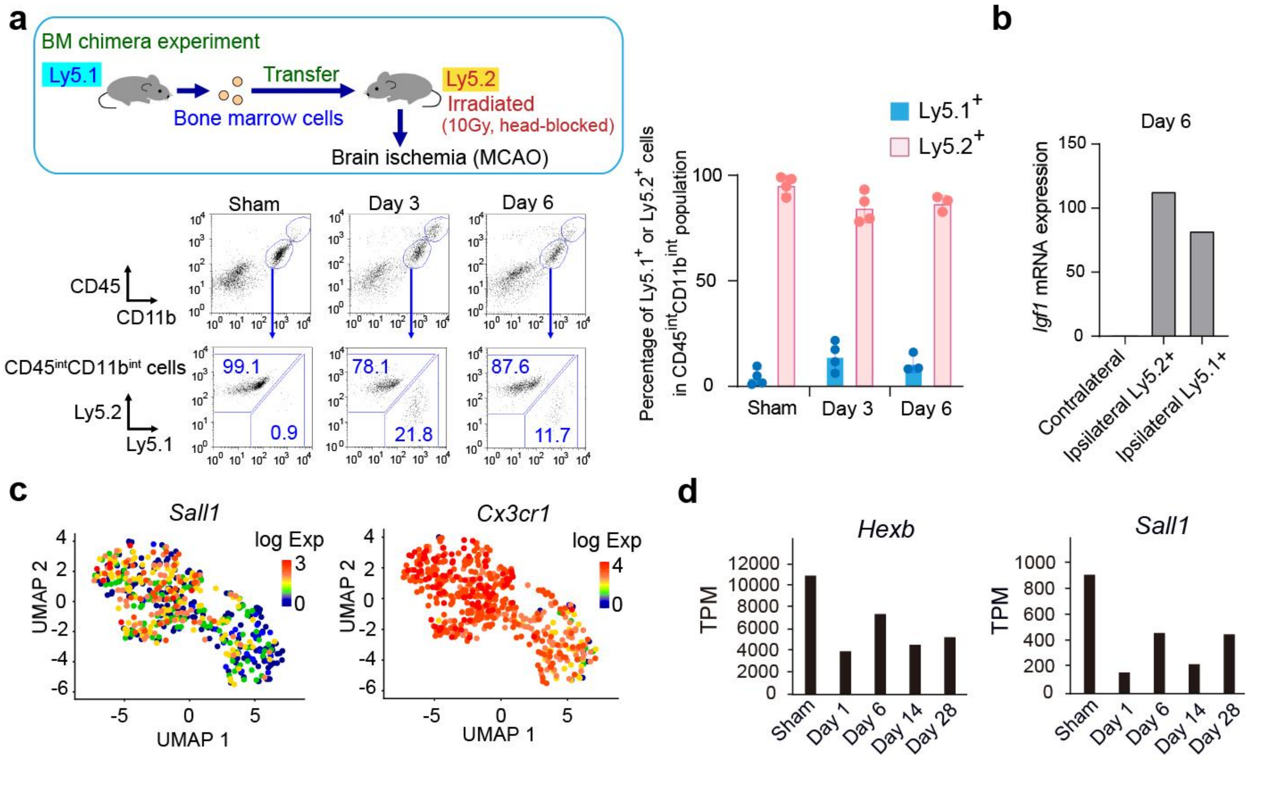


**Extended Data Figure 3. Analysis of $CD45^{int}CD11b^{int}$ cells after ischemic stroke onset.** (**a**) Left panel: FACS analysis of $CD45^{int}CD11b^{int}$ cells collected from sham-operated or post-ischemic brains of $Ly5.1^{+}$ bone marrow-transferred $Ly5.2^{+}$ mice. Right panel: the percentage of $Ly5.1^{+}$ or $Ly5.2^{+}$ cells in $CD45^{int}CD11b^{int}$ cells of $Ly5.1^{+}$ bone marrow-transferred $Ly5.2^{+}$ mice at each time point after stroke onset. (**b**) Relative *Igf1* mRNA expression levels in each $CD45^{int}CD11b^{int}$ cell collected from the contralateral or ipsilateral hemisphere on day 6 after stroke onset. Cells were collected from at least 6 mice after stroke onset. (**c**) Gene expression heatmaps of post-ischemic day 14 $CD45^{int}CD11b^{int}$ cells. (**d**) Time-dependent change of *Hexb* or *Sall1* mRNA expression levels detected by RNA-seq analysis in $CD45^{int}CD11b^{int}$ cells collected from at least 10 mice after stroke onset. Error bars represent the mean ± standard error of the mean (SEM).

**a**

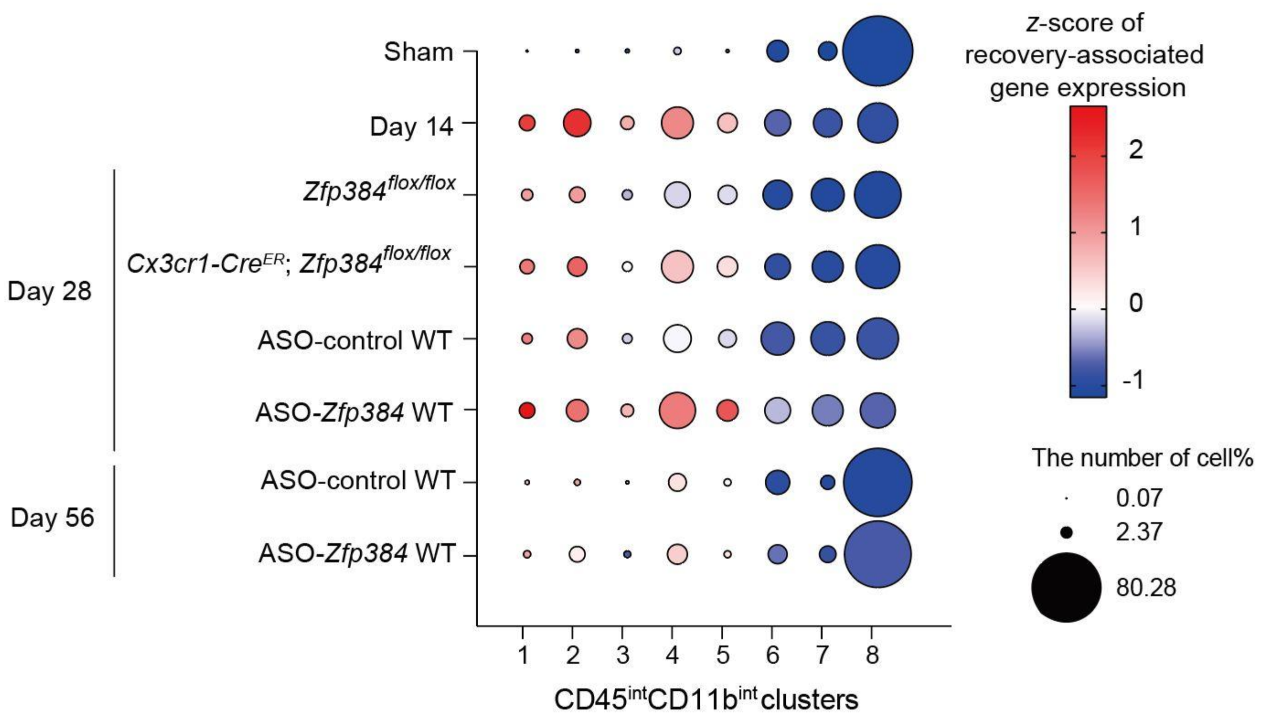


**Extended Data Figure 4. The time-dependent changes of CD45$^{int}$CD11b$^{int}$ clusters before and after ischemic stroke onset.** Bubble heatmap chart showing the read ratio of recovery phase–associated genes. The size of the circles indicates the percentage of each cluster in CD45$^{int}$CD11b$^{int}$ cells.

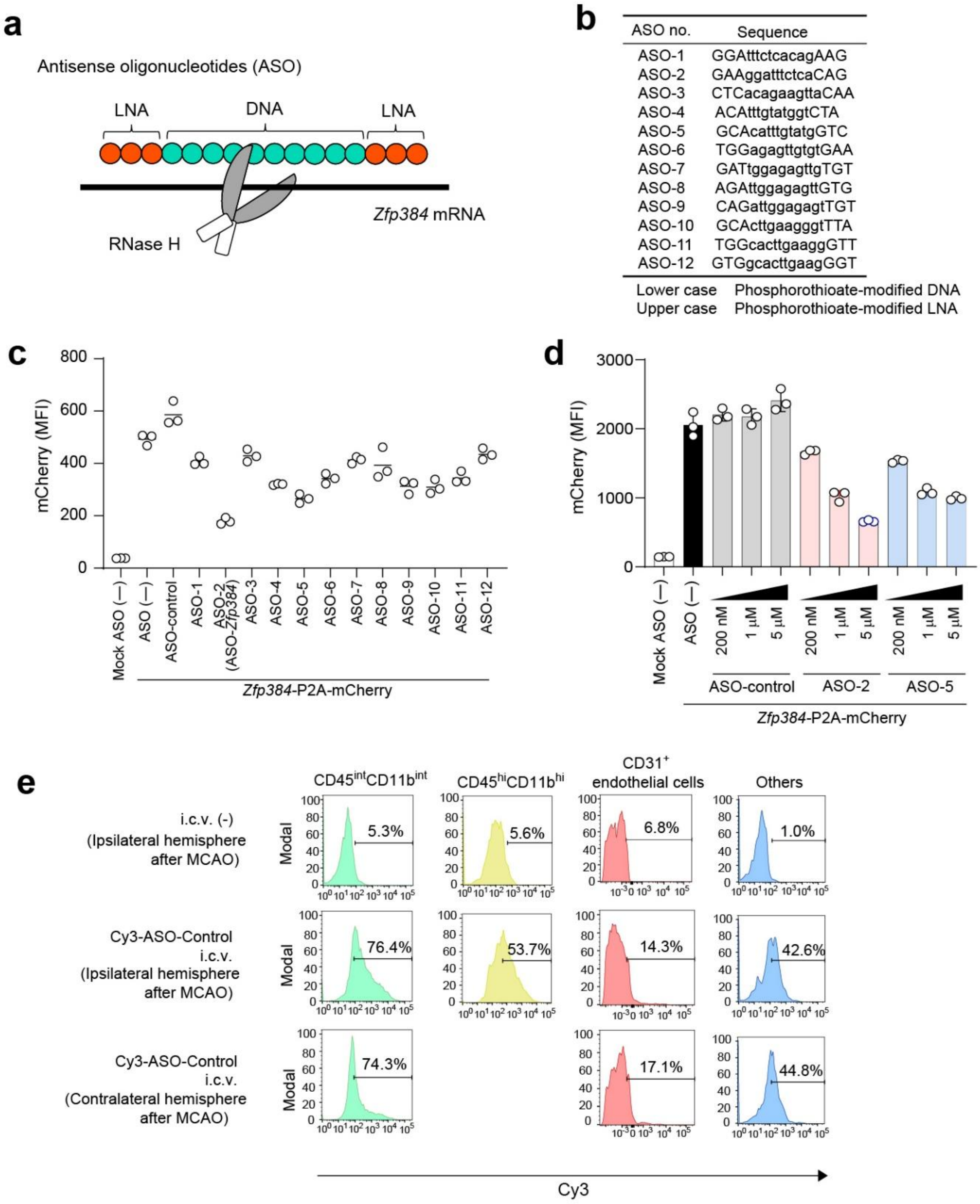


**Extended Data Figure 5. Development of antisense oligonucleotides (ASOs) against *Zfp384*.** (**a**) Scheme of ASOs designed for *Zfp384*. (**b**) The list of the ASO sequences designed in this study. (**c**) Mean fluorescence intensity (MFI) of cells showing mCherry fluorescence in mock-transduced or Zfp384-P2A-mCherry overexpressing microglial cell line. ASO-mediated suppression of *Zfp384*-P2A-*mCherry* expression was evaluated by FACS. (**d**) Dose-dependent suppression of *Zfp384*-P2A-*mCherry* expression in ASO-treated microglial cell line. (**e**) Confirmation of the internalization of Cy3-conjugated ASO in each brain cell type when ASO was administered into the lateral ventricle after the induction of ischemic stroke.

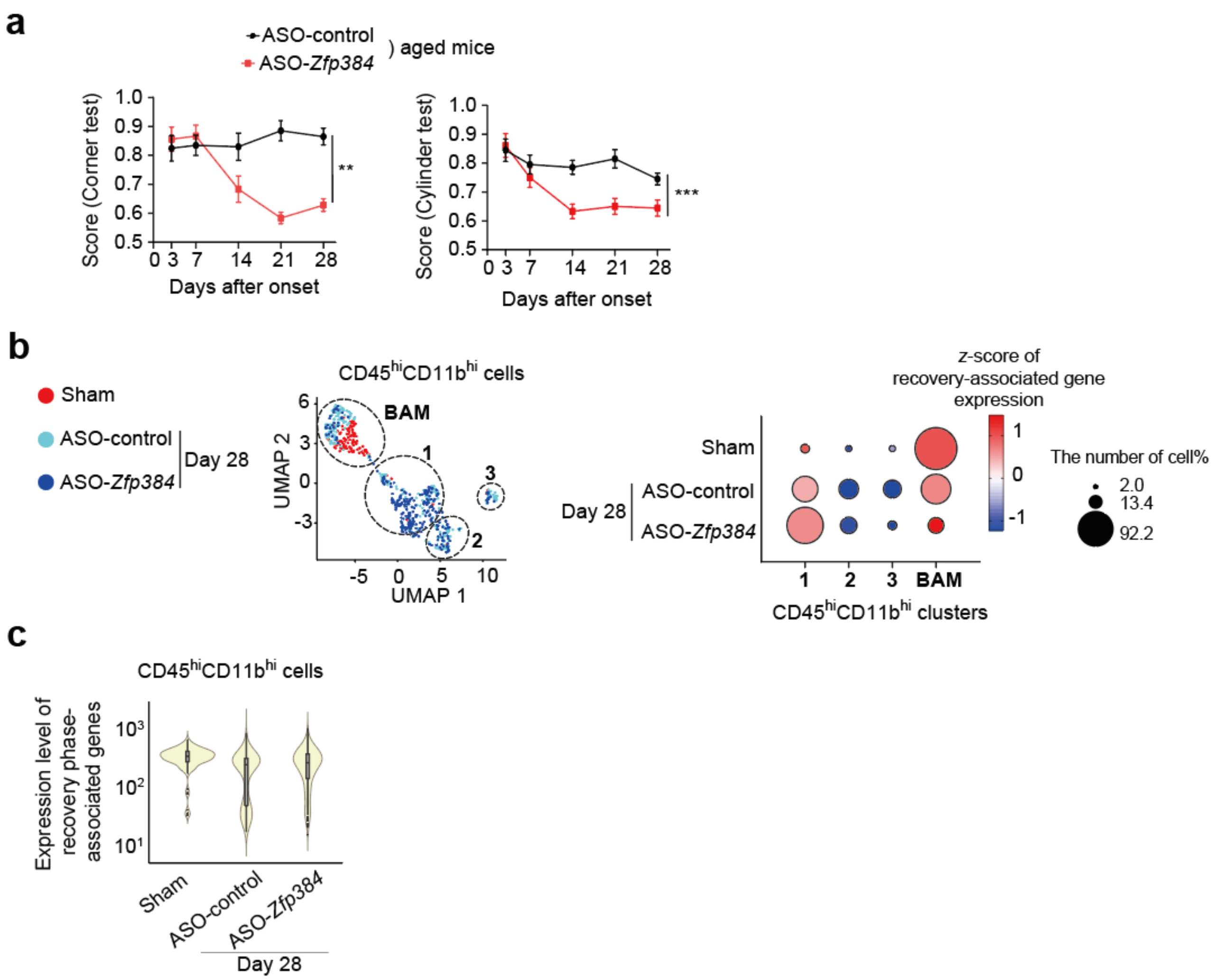


**Extended Data Figure 6. The effects of ASO-*Zfp384* in aged mice and CD45$^{hi}$CD11b$^{hi}$ cells.** (**a**) Neurological deficits after stroke onset in aged mice administered ASO-control or ASO-*Zfp384* (n = 10 for ASO-control, 12 for ASO-*Zfp384*). ASOs were administered 8 and 22 days after stroke onset. (**b**) UMAP of CD45$^{hi}$CD11b$^{hi}$ cells (left panel) and bubble heatmap chart showing the read ratio of recovery phase-associated genes (right panel). The circles' size indicates each cluster's percentage in CD45$^{hi}$CD11b$^{hi}$ cells. BAM: border-associated macrophage which was detected by *Ms4a7* and *Mrc1* expression (**c**) Comparison of total standardized read counts of recovery phase–associated genes in all CD45$^{hi}$CD11b$^{hi}$ cells at each time point. **$p < 0.01$, ***$p < 0.001$ vs. ASO-control (**a**) (two-way ANOVA with Tukey's test [**a**]). Error bars represent the mean ± standard error of the mean (SEM).

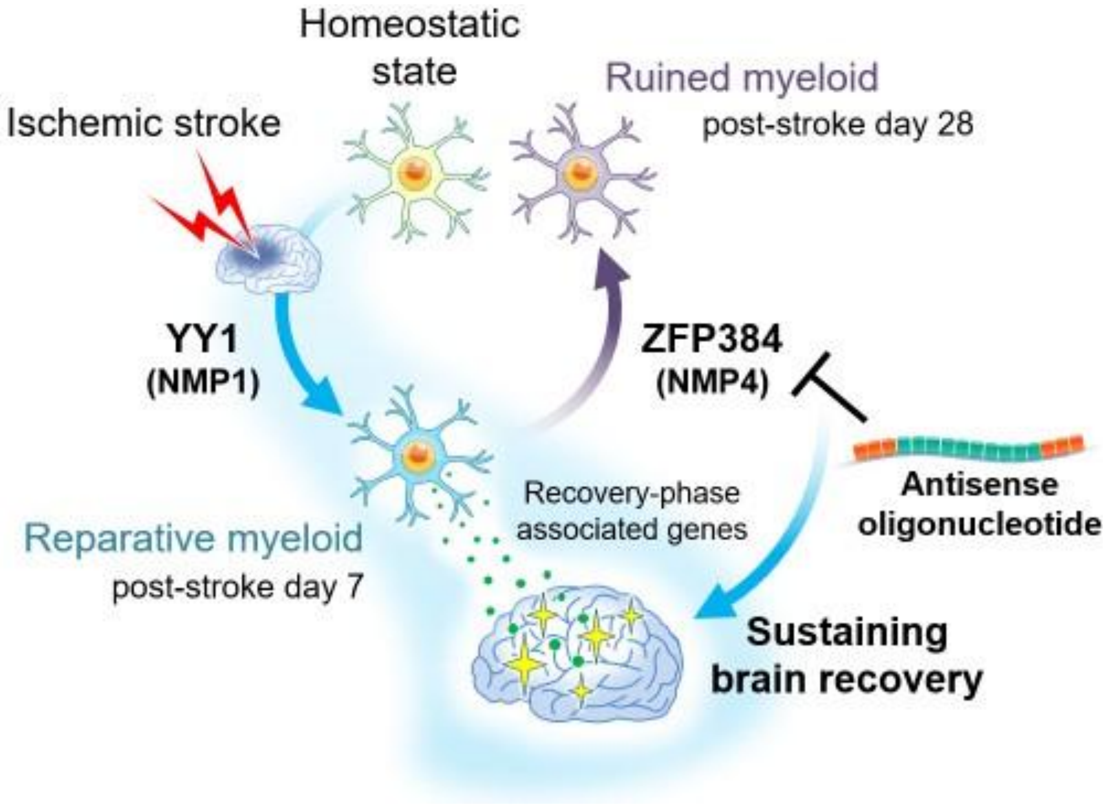


**Extended Data Figure 7. Summarized figure.**

Almost spontaneous brain recovery will be terminated a few months after brain injury, although the cellular fate of reparative microglia has not been clarified. Ischemic stroke induces reparative myeloid cells around 7 days after stroke onset, expressing recovery phase–associated genes through the transcriptional regulation by YY1 (nuclear matrix protein 1: NMP1). ZFP384 (NMP4) diminishes a broad range of neural repair functions in myeloid cells to reach a ruined state by abrogating YY1-mediated expression of recovery phase–associated genes. Antisense oligonucleotide against *Zfp384* prolongs their reparative functions to sustain brain recovery and improve functional prognosis.

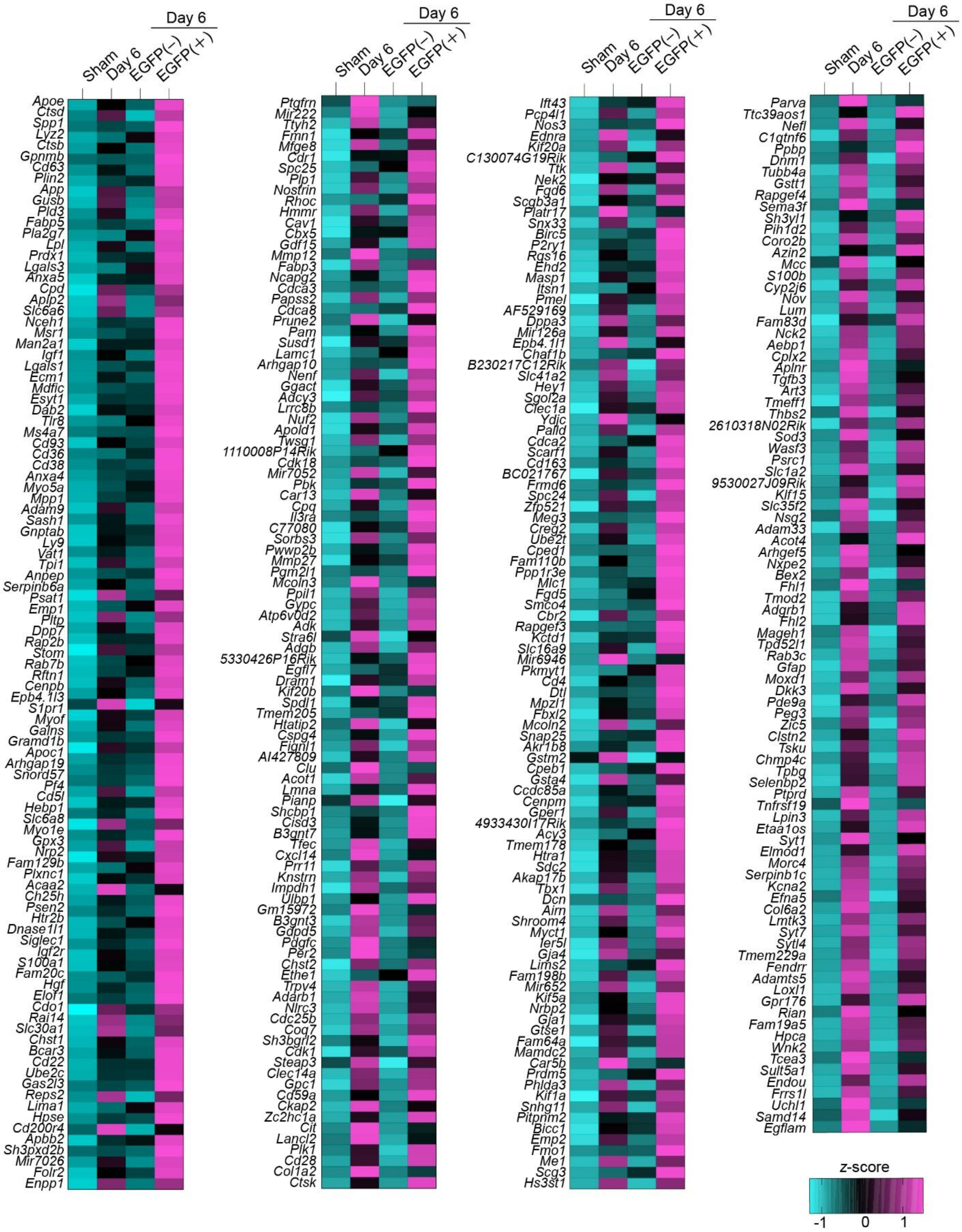


**Extended Data Table 1. The list of 391 recovery phase–associated genes.**

**a**

| | *Zfp384*$^{flox/flox}$ | *Cx3cr1-Cre*$^{ER}$; *Zfp384*$^{flox/flox}$ |
|---|---|---|
| MABP (mmHg) | 85 ± 4 | 84 ± 5 |
| pH | 7.40 ± 0.02 | 7.39 ± 0.02 |
| $PaO_2$ (mmHg) | 141 ± 7 | 139 ± 3 |
| $PaCO_2$ (mmHg) | 26.4 ± 2.3 | 25.5 ± 1.7 |
| Hematocrit (%) | 35 ± 1 | 36 ± 1 |
| Glucose (mg/dl) | 242 ± 14 | 263 ± 8 |

**b**

| CBF reduction (%) | before CCA occulusion | after CCA occulusion | after MCA occulusion | N | Exclusion (N) Death | CBF |
|---|---|---|---|---|---|---|
| *Zfp384*$^{flox/flox}$ | 100 | 79.4 ± 2.6 | 29.8 ± 1.5 | 14 | 0 | 0 |
| *Cx3cr1-Cre*$^{ER}$; *Zfp384*$^{flox/flox}$ | 100 | 82.1 ± 2.3 | 29.7 ± 1.5 | 15 | 0 | 0 |

**c**

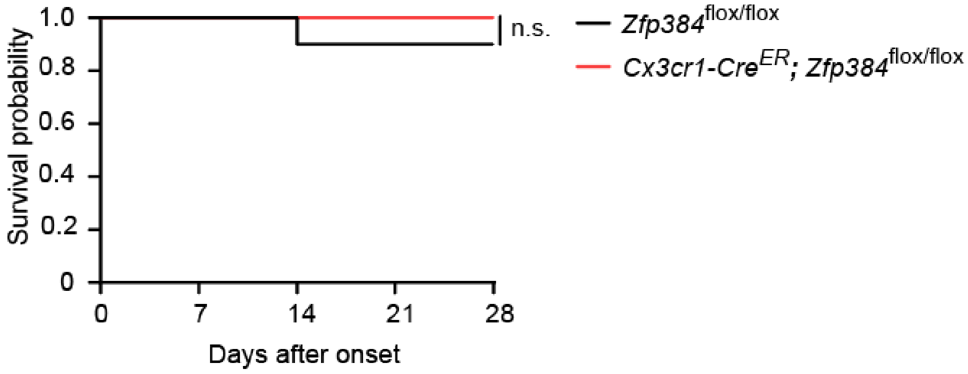


**Extended Data Table 2. Results of physiological data, cerebral blood flow, and survival in *Zfp384*$^{flox/flox}$ and *Cx3cr1-Cre*$^{ER}$; *Zfp384*$^{flox/flox}$ mice.** (**a**) Physiological data from each *Zfp384*$^{flox/flox}$ mice. (**b**) Changes in cerebral blood flow before and after the induction of brain ischemia. The number of mice excluded upon death or non-satisfactory reduction of CBF during ischemia was shown. (**c**) Survival rate after stroke onset in *Zfp384*$^{flox/flox}$ and *Cx3cr1-Cre*$^{ER}$; *Zfp384*$^{flox/flox}$ mice. n.s.: not significant. (**a**,**b**,**c**)There was no significant difference in all parameters shown in this table.

**a**

| CBF reduction (%) | before CCA occulusion | after CCA occulusion | after MCA occulusion | N | Exclusion (N) Death | CBF |
|---|---|---|---|---|---|---|
| ASO-control i.c.v. → WT | 100 | 81.3 ± 2.6 | 29.1 ± 1.0 | 10 | 0 | 0 |
| ASO-*Zfp384* i.c.v. → WT | 100 | 81.9 ± 3.0 | 29.4 ± 0.6 | 10 | 0 | 0 |

**b**

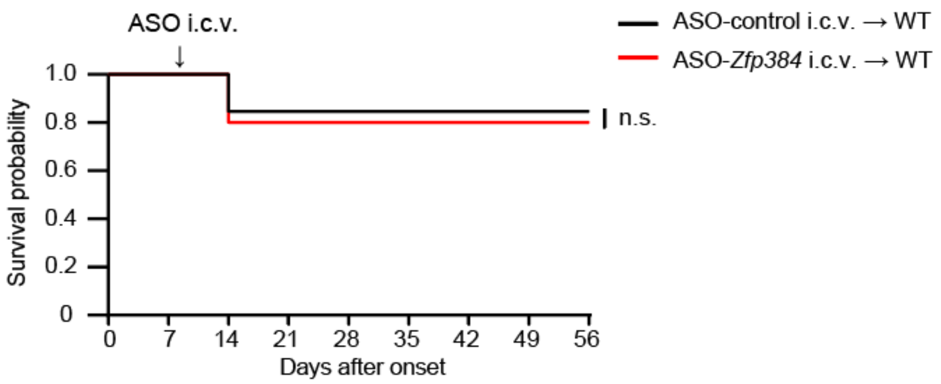


**c**

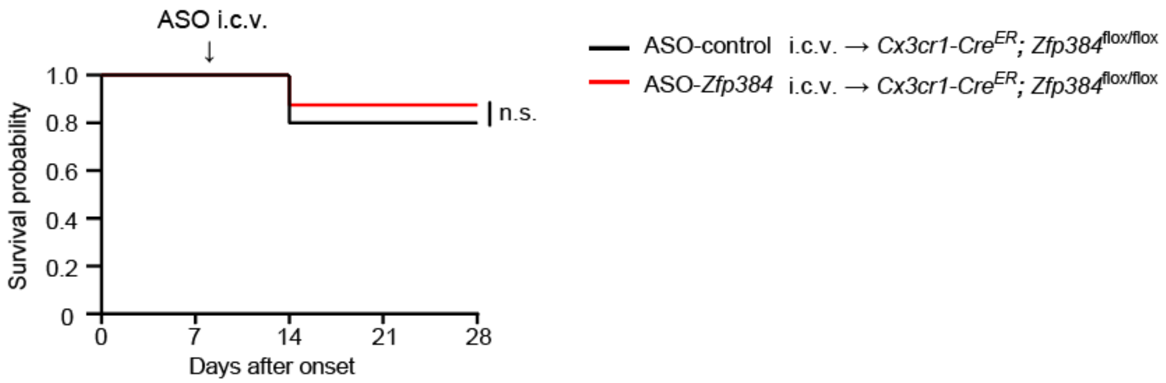


**Extended Data Table 3. Cerebral blood flow and survival in ASO-administered mice.** (**a**) Changes in cerebral blood flow before and after the induction of brain ischemia. There was no significance between the two groups. The number of mice excluded upon death or non-satisfactory reduction of CBF during ischemia was shown. (**b**,**c**) Survival rate after stroke onset in ASO-administered WT mice (**b**) or ASO-administered *Cx3cr1-Cre*$^{ER}$; *Zfp384*$^{flox/flox}$ mice (**c**). n.s.: not significant.
